# The fission yeast cell size control system integrates pathways measuring cell surface area, volume, and time

**DOI:** 10.1101/2022.11.17.516946

**Authors:** Kristi E. Miller, Cesar Vargas-Garcia, Abhyudai Singh, James B. Moseley

## Abstract

Eukaryotic cells tightly control their size, but the relevant aspect of size is unknown in most cases. Fission yeast divide at a threshold cell surface area due in part to the protein kinase Cdr2. We find that fission yeast cells only divide by surface area under a size threshold but shift to volume-based divisions when they reach a larger size. The size threshold for changing from surface area to volume-based control is set by ploidy. Within this size control system, we identified the mitotic activator Cdc25 as a volume-based sizer molecule, while the mitotic cyclin Cdc13 accumulates as a timer. We propose an integrated model for cell size control based on multiple signaling pathways that report on distinct aspects of cell size and growth, including cell surface area (Cdr2), cell volume (Cdc25), and time (Cdc13). Combined modeling and experiments show how this system can generate both sizer and adder-like properties.

**HIGHLIGHTS:**

- Fission yeast use surface area or volume-based cell size control depending on overall size and ploidy
- Mitotic activator Cdc25 exhibits properties of a volume-based sizer molecule
- Mitotic cyclin Cdc13 accumulates in the nucleus dependent on time, not size
- Combined modeling and experiments identify conditions for sizer versus adder behavior

## INTRODUCTION

Cell size varies widely among different cell types, but most cells control their size within an optimal window for proper function (Amodeo & Skotheim, 2016; Dolznig et al., 2004; Ginzberg et al., 2015). Three common strategies for cell size control are timers, adders, and sizers (Jun & Taheri-Araghi, 2015; Facchetti et al., 2017). Timers act by limiting cell cycle stages to defined periods of time, as often observed in early stages of development (Donnan & John, 1983; Wang et al., 2000). Adders measure the amount of growth or mass added to a cell during a defined cell cycle stage, as seen in bacteria and mammalian cells (Campos et al., 2014; Taheri-Araghi et al., 2015; Varsano et al., 2017). Sizers work by delaying cell cycle transitions until cells reach a critical size threshold. Sizers have been identified in bacteria, yeast, and animal cells, and they imply the existence of size measuring systems that signal to core cell cycle machinery (Fantes, 1977; Schmoller et al., 2015; Sveiczer et al., 1996; Talia et al., 2007; Turner et al., 2012; Wallden et al., 2016; Xie & Skotheim, 2020). As cells grow, they increase in volume, surface area, mass, and more. For most cell types that exhibit sizer behavior, we do not know which aspect of size is measured, or whether different molecular pathways measure distinct aspects of cell size. Many previous studies have focused on size-dependent changes in an individual protein or pathway, leaving open questions regarding how multiple pathways could converge to generate a robust and dynamic size control system.

The fission yeast *Schizosaccharomyces pombe* is a strong model system for studies on sizer behavior. These rod-shaped cells grow by linear extension while maintaining a constant cell width (Mitchison & Nurse, 1985). Fission yeast cells adjust the amount of growth per cell cycle to compensate for variations in size at birth, such that all cells enter mitosis and divide at a defined and reproducible size (Fantes, 1977; Sveiczer et al., 1996). Historically, cell length has been used as a proxy for cell size in fission yeast studies, but recent work using mutants with altered widths has shown that fission yeast cells divide at a defined surface area, as opposed to volume or length (Pan et al., 2014; Facchetti et al., 2019). This finding raises the question of how cells measure their surface area, and whether other aspects of cell growth and geometry contribute to the overall size control system.

A set of conserved regulatory proteins controls mitotic entry and thereby cell size in fission yeast (Figure 1A). The cyclin-dependent kinase Cdk1 (also called Cdc2 in *S. pombe*) drives mitotic entry in complex with the B-type cyclin Cdc13 (Harashima et al., 2013). The protein kinase Wee1 inhibits Cdk1 in small cells by phosphorylating a conserved tyrosine residue (Russell & Nurse, 1987; Gould & Nurse, 1989). This inhibition is reversed by the protein phosphatase Cdc25 to promote mitotic entry (Russell & Nurse, 1986; Moreno et al., 1990; Gautier et al., 1991; Kumagai & Dunphy, 1991; Strausfeld et al., 1991).

**Figure 1.**
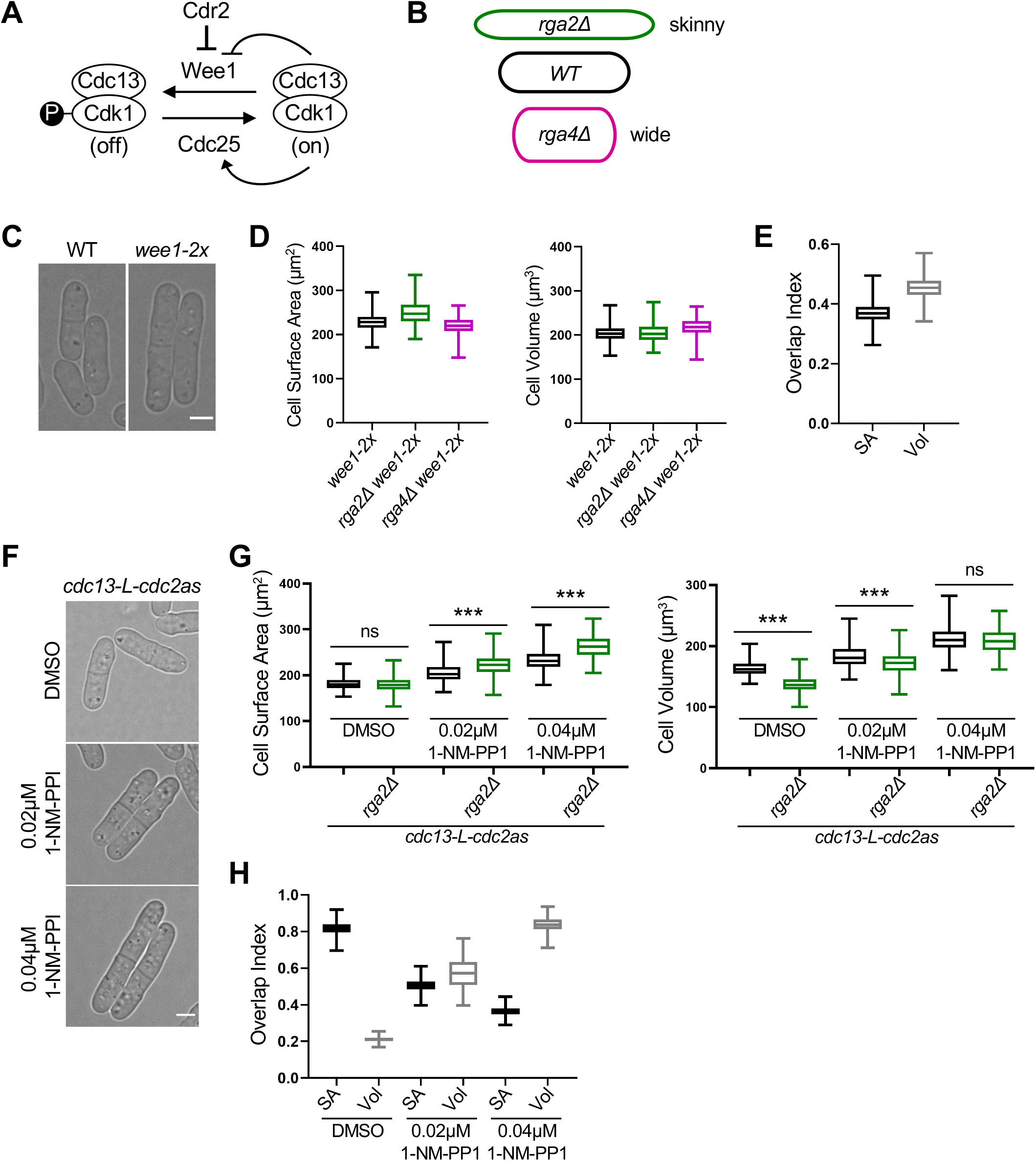
Large cells shift to volume-based division. (A) Schematic of the pathway regulating mitotic entry and cell size. (B) Cartoon of cell width mutants used to uncouple cell surface area and volume. The colors apply to subsequent graphs in all figures. (C) Brightfield images of wild type (WT) and larger *wee1-2x* cells. (D) Size of dividing cells for indicated strains plotted by surface area (left) or volume (right). (E) Overlap index analysis for data in panel D. SA, surface area. Vol, volume. (F) Brightfield images of *cdc13-L-cdc2as* treated with the indicated concentrations of 1-NM-PP1 or DMSO control. (G) Size of dividing cells for indicated strains plotted by surface area (left) or volume (right). ns, not significant; ***p<0.0001. (H) Overlap analysis for data in panel G. Graphs shows median as a line, quartiles, max, and min. Bars, 4μm.

Within this conserved regulatory module, three mitotic activators have been heavily studied as potential size-measuring systems. First, the protein kinase Cdr2 controls cell size-dependent inhibition of Wee1 at a series of cortical multiprotein structures called “nodes” (Young & Fantes, 1987; Breeding et al., 1998; Kanoh & Russell, 1998; Allard et al., 2018). The cortical density of nodes scales with cell surface area and could act as a surface area measurement system (Facchetti et al., 2019; Pan et al., 2014). Indeed, loss of Cdr2 causes cells to divide at an increased size and at a reproducible volume, as opposed to surface area (Facchetti et al., 2019; Opalko et al., 2022). However, it is not known if this shift from surface area-based to volume-based size control reflects changes in overall cell size or loss of Cdr2 signaling. A second potential size-measuring system in fission yeast is accumulation of the cyclin Cdc13. Unlike most proteins, the cellular concentration of Cdc13 increases as cells grow, a phenomenon known as ‘superscaling.’ Cdc13 accumulates in the nucleus, where it can complex with Cdk1, and this superscaling has been proposed to act as a primary sensor for the mitotic size control system (Patterson et al., 2019). A third possible size measuring system is superscaling of Cdc25 phosphatase (Moreno et al., 1990; Keifenheim et al., 2017). A recent study found that Cdc13 and Cdc25 are the only known mitotic regulatory proteins that increase in concentration as cells grow (Curran et al., 2022). However, it has not been known what aspect of cell size or growth drives accumulation of these mitotic activators.

Here, we studied connections between cell geometry, scaling of mitotic activators, and size control. We show that Cdc13/cyclin accumulates as a molecular timer, as opposed to a sizer. In contrast, Cdc25 accumulates specifically with cell volume, not with surface area or time. Thus, fission yeast cells have distinct pathways related to cell surface area (Cdr2-Wee1), cell volume (Cdc25), and time (Cdc13/cyclin). By altering cell size and mitotic signaling with previously characterized mutations, we identified conditions where fission yeast cells stop dividing by surface area-based size control and shift to other modes including volume-based size control and adder-like behavior. Our results redefine the fission yeast size control system as a network that integrates multiple pathways, each of which monitors distinct aspects of cell size and growth. Our model has strong implications for mechanisms of size control in other cell types and organisms.

## RESULTS

### Large cells shift to volume-based size control

We started our study by asking how changes to overall cell size impact surface area-based size control. We used wide (*rga4*Δ) and skinny (*rga2*Δ) mutants to uncouple the scaling of cell surface area, volume, and length, similar to previous work (Figure 1B) (Pan et al., 2014; Facchetti et al., 2019; Opalko et al., 2022). We designed a semi-automated image analysis pipeline to measure cell geometry (Figure S1A), which confirmed that wild type cells divide at a constant surface area while *cdr2*Δ cells divide at a constant volume (Figures S1B-S1E). We also generated an “overlap index” to quantify whether sizes were more related by surface area or by volume (see Methods). This index confirmed the shift to volume-based divisions for *cdr2*Δ mutant cells (Figure S1E).

*cdr2*Δ cells divide at an increased size compared to wild type, so the shift to volume-based divisions could reflect the change in cell size or alternatively the loss of Cdr2 signaling. To distinguish between these possibilities, we tested other large cell size mutants. *wee1-2x* cells express an extra copy of *wee1+*. These cells divided larger than wild type but similar to *cdr2*Δ mutants (Figures 1C and S1F). Combining *wee1-2x* with wide (*rga4*Δ) or skinny (*rga2*Δ) mutants revealed that these enlarged cells also divide at a constant volume, as opposed to surface area (Figures 1D and 1E). This result suggests that large cells switch to volume-based division even with Cdr2 present. As an additional test, we used cells expressing *cdc13-L-cdc2as*, in which the activity of Cdk1 (i.e. Cdc2) is inhibited by 1-NM-PP1 (Coudreuse & Nurse, 2010). We controlled the size of this strain by adding two different concentrations of 1-NM-PP1, leading to concentration-dependent increases in cell size compared to a DMSO control (Figure S1F). Interestingly, we observed a 1-NM-PP1 concentration-dependent shift towards volume-based division as cell size increased, which was validated by the overlap index (Figures 1G and 1H). We conclude that cells shift from surface area-based division to volume-based division above a certain size threshold.

The shift of *cdr2*Δ mutants to volume-based division has been suggested to result from loss of surface area-sensing by the Cdr2 pathway (Facchetti et al., 2019), but we observed a similar shift for large cells with an intact Cdr2 pathway. These results raise the possibility that the shift is due to increased size, as opposed to a specific role for Cdr2. These possibilities can be distinguished by reducing the size of *cdr2*Δ mutants. Therefore, we combined *cdr2*Δ with the previously characterized *ppa2*Δ and *zfs1*Δ mutations, which additively reduce cell size (Kinoshita et al., 1993; Navarro & Nurse, 2012). The resulting triple mutant *cdr2*Δ *ppa2*Δ *zfs1*Δ divided at the same size as wild type cells (Figures 2A and 2B). By combining *cdr2*Δ*ppa2*Δ *zfs1*Δ with wide (*rga4*Δ) and skinny (*rga2*Δ) mutants, we found that reducing the size of *cdr2*Δ mutants shifted cells back to surface area-based divisions (Figure 2C). We note that the overlap index for surface area in this experiment was reduced compared to wild type cells, raising the possibility of altered size control for this strain (Figure 2D). Nonetheless, the loss of surface area-based divisions in *cdr2*Δ mutants is largely suppressed by reducing cell size, supporting a model where overall size determines whether cells divide by surface area or by volume. This result is not inconsistent with Cdr2 acting as a surface area sensor, but suggests that Cdr2 acts together with additional, redundant surface area measuring systems.

**Figure 2.**
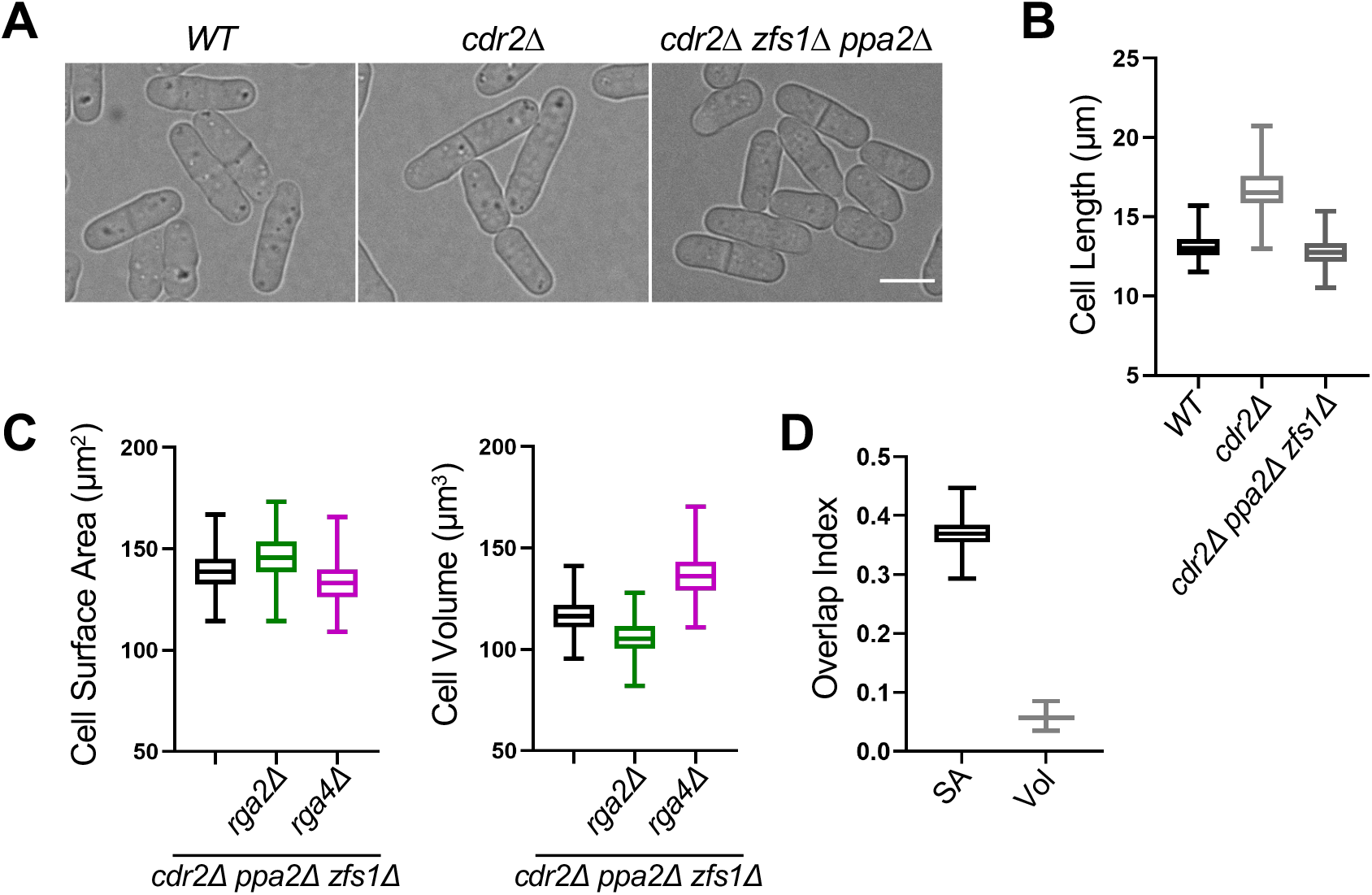
Reducing the size of *cdr2*Δ cells restores surface area-based divisions. (A) Brightfield images of WT, *cdr2*Δ, and *cdr2*Δ *zfs1*Δ *ppa2*Δ cells. Bar, 4μm. (B) Cell length at division (μm) of indicated cell types. (C) Size of dividing cells for indicated strains plotted by surface area (left) or volume (right). (D) Overlap index analysis for data in panel C. SA, surface area. Vol, volume. Graphs shows median as a line, quartiles, max, and min.

Next, we tested if loss of surface area-based divisions is specific to increased cell size or also occurs when cells divide too small. We examined the small size mutant *ppa2*Δ, previously shown to divide at a smaller size than wild type cells (Figures 3A and S1F). *ppa2*+ encodes one of two partially redundant catalytic subunits of the PP2A phosphatase, which counteracts Cdk1 activity (Kinoshita et al., 1993). We found that *ppa2*Δ, *ppa2*Δ *rga2*Δ, and *ppa2*Δ *rga4*Δ all divided at the same surface area, but not at the same volume (Figures 3B and 3C). Therefore, reduced cell size does not cause a shift to volume-based divisions. Rather, the shift to volume-based divisions is a specific property of larger cell size.

**Figure 3.**
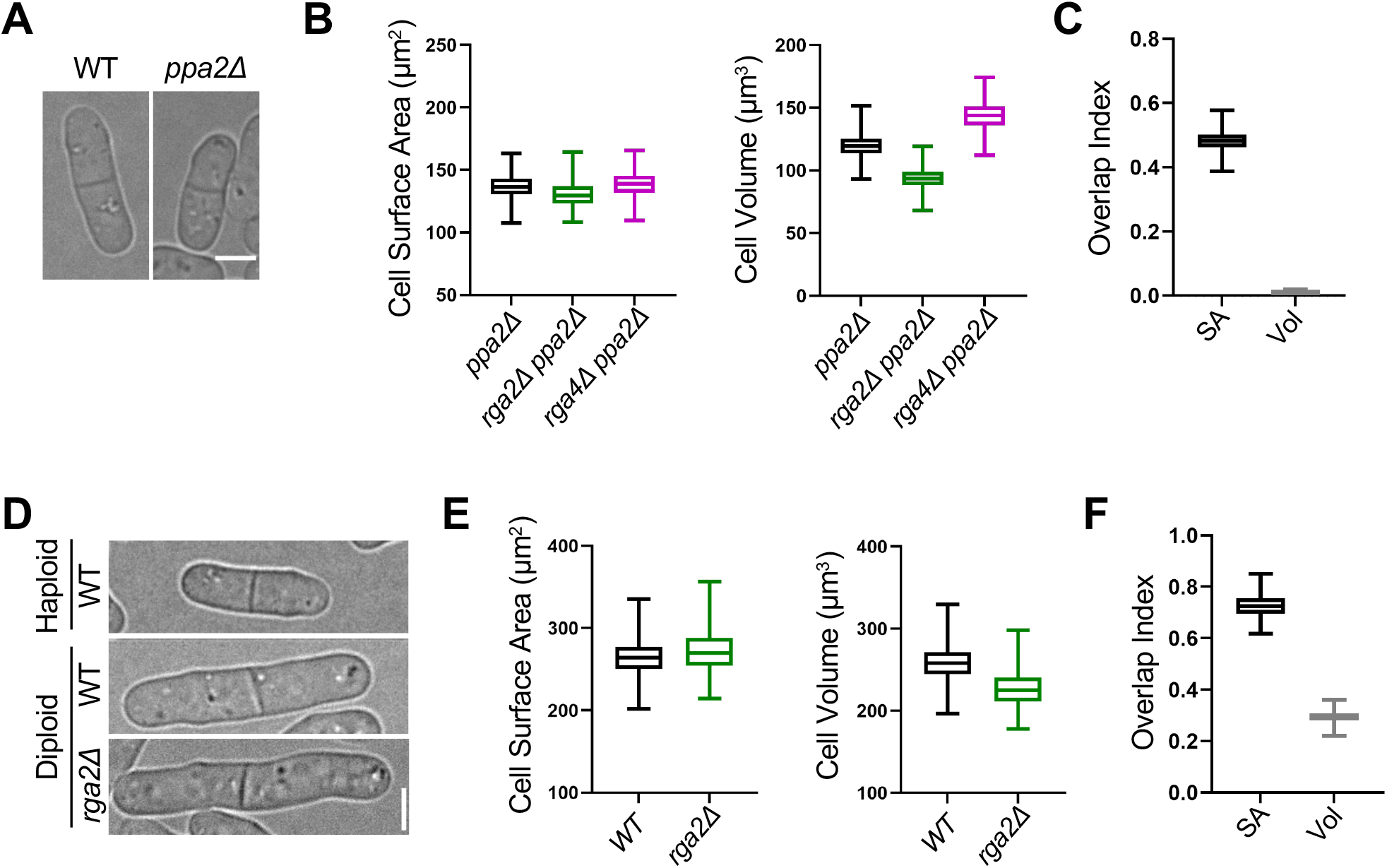
Small size mutants and diploid cells retain surface area-based divisions. (A) Brightfield images of WT and small *ppa2*Δ cells. (B) Size of dividing cells for indicated strains plotted by surface area (left) or volume (right). (C) Overlap index analysis for data in panel B. SA, surface area. Vol, volume. (D) Brightfield images of indicated strains. (E) Size of dividing cells for indicated strains plotted by surface area (left) or volume (right). (F) Overlap index analysis for data in panel E. Graphs shows median as a line, quartiles, max, and min. Bars, 4μm.

Increased cell size has also been observed to scale with increased ploidy for many cell types and organisms (Figure 3D) (Amodeo & Skotheim, 2016). Our results were obtained in haploid cells, so we hypothesized that genome content could limit the ability of large cells to maintain surface area-based divisions. In support of this model, previous studies have shown that genome content limits transcription in large fission yeast cells (Zhurinsky et al., 2010), and the cytoplasm of budding yeast cells becomes diluted with increased size (Schmoller et al., 2015; Neurohr et al., 2019). We found that wild type and *rga2*Δ diploid fission yeast cells divide at an overlapping surface area as opposed to volume, despite increased overall size (Figures 3E and 3F). Therefore, diploid cells use surface area-based divisions similar to haploid cells, despite their increased overall size. This result indicates that the size threshold for shifting from surface area-based to volume-based divisions is set by ploidy.

### Cdc25 nuclear accumulation monitors cell volume

We next sought to identify molecular mechanisms that act in cell surface area-based and volume-based size control. Previous work has shown that Cdr2 localization patterns scale with cell surface area and may contribute to cell division at a threshold surface area (Pan et al., 2014; Facchetti et al., 2019), but no volume-based sizer molecules have been identified. Here, we focused on the Cdk1-activating phosphatase Cdc25 and the B-type mitotic cyclin Cdc13. The concentrations of both Cdc25 and Cdc13 increase as cells grow (Moreno et al., 1990; Keifenheim et al., 2017; Patterson et al., 2019; Curran et al., 2022), but it has not been known what aspect of size and/or growth this accumulation reflects. Both Cdc25 and Cdc13 accumulate in the nucleus where Cdk1 activation likely occurs, so we focused on their nuclear concentrations. It is also important to note that the size of the nucleus scales tightly with overall cell size (Cantwell & Nurse, 2019; Neumann & Nurse, 2007). As a starting point, we used functional Cdc25-mNG (mNeonGreen) expressed from the endogenous promoter and chromosomal locus. For imaging and segmenting the nucleus, cells also expressed a nuclear localization sequence fused to blue fluorescent protein (BFP-NLS). Cdc25-mNG and BFP-NLS were imaged by confocal microscopy in wild type, wide (*rga4Δ*), and skinny (*rga2*Δ) cells. Brightfield imaging of cells was performed to obtain cell dimensions and to segment cells (Figure S1A). To rapidly measure cell geometry and fluorescent protein intensity in thousands of cells, we developed a semi-automated ImageJ/MATLAB image analysis pipeline (Figure S1A). With this pipeline, we confirmed that the whole-cell concentration of both Cdc25 and Cdc13 increases with cell size (Figures S3F and S4F).

We plotted the mean Cdc25-mNG intensity in the nucleus as a function of cell surface area or cell volume. We observed strong overlap for Cdc25-mNG nuclear concentration in wild type, *rga4Δ*, and *rga2*Δ cells when plotted by volume but not by surface area (Figure 4A). This result suggests that the nuclear accumulation of Cdc25 scales with cell volume, not with cell surface area. This conclusion was strengthened when we excluded large, potentially mitotic cells (Figure S3A), and compared groups of cells binned by size (Figures 4B and S3D). We also reached the same conclusion quantifying either mean intensity or sum intensity of Cdc25-mNG in the nucleus (Figures S3A-C). In all analyses, Cdc25 nuclear concentration scaled closely with cell volume but not surface area. We conclude that Cdc25 nuclear concentration is a volume-dependent input to the cell size control network.

**Figure 4.**
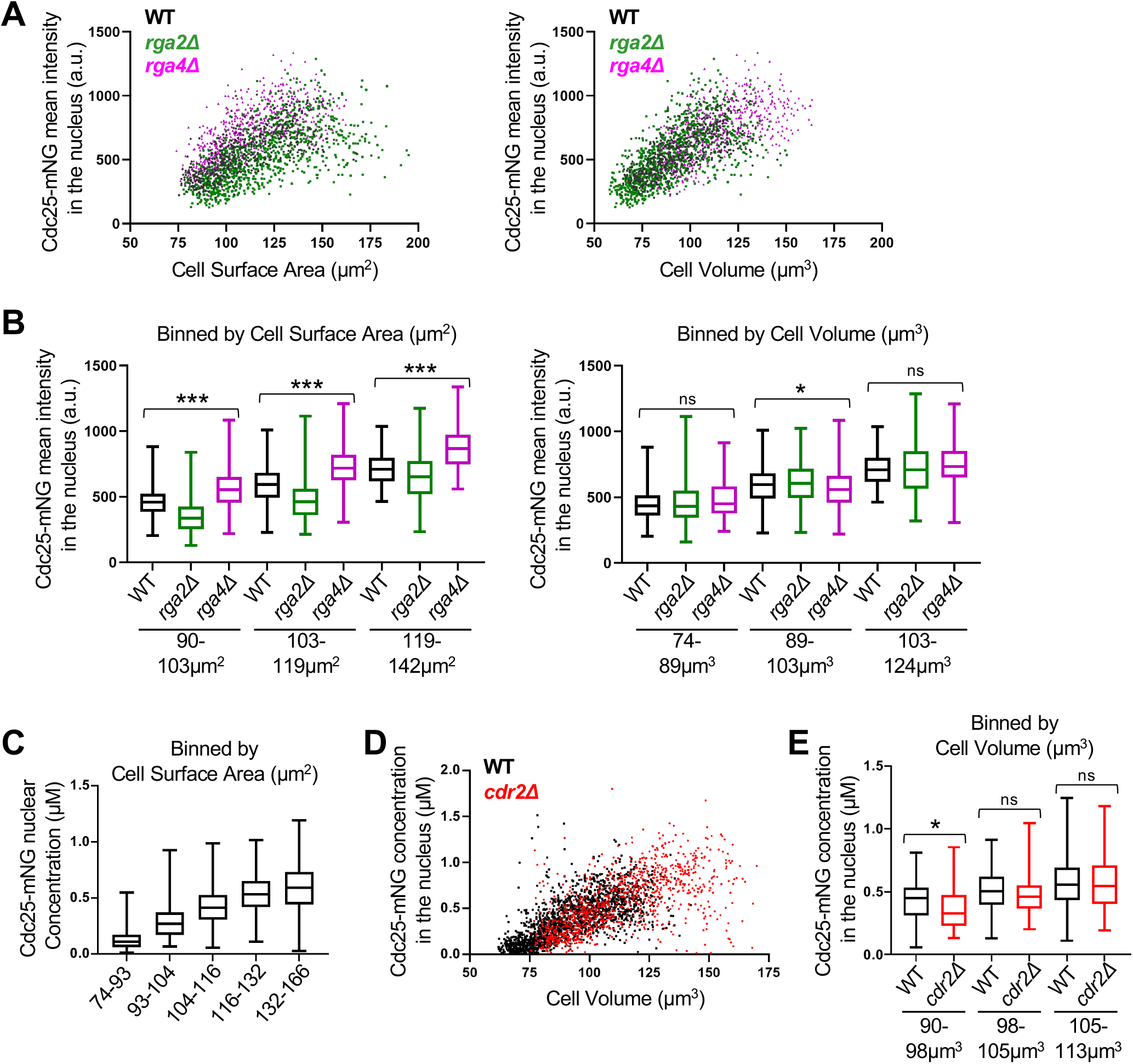
Cdc25 accumulation in the nucleus scales with cell volume. (A) Cdc25-mNG nuclear mean intensity in WT, *rga2*Δ, and *rga4*Δ single cells plotted either by cell surface area (left) or by cell volume (right) (WT n=502, *rga2*Δ n=867, and *rga4*Δ n=684). The same cells are plotted in the two graphs. (B) Cdc25 mean intensity in the nucleus for WT, *rga2*Δ, and *rga4*Δ cells binned by cell surface area (left) or volume (right). ***p<0.0001; *p<0.03; ns, not significant. n>90 for each strain per bin. Reported p value is the most significant value from pairwise comparisons within each group. (C) The concentration (μM) of nuclear Cdc25-mNG increases with cell size. Cells are binned by cell surface area (n>250 for each size bin). (D) Cdc25-mNG nuclear concentration (μM) in WT and *cdr2*Δ cells plotted by their cell volume (WT n=1,801; *cdr2*Δ n=1,037). (E) WT and *cdr2*Δ cells of the same volume have similar nuclear concentration (μM) of Cdc25 (n>110 per strain). Graph shows median as a line, quartiles, max, and min. *p=0.004.

We performed three additional experiments to extend this conclusion. First, we calibrated our confocal microscope using proteins of known concentration to measure the molar concentration of Cdc25-mNG in the nucleus (Wu & Pollard, 2005) (Figure S3E). In small cells, Cdc25 was present in the nucleus at levels below 100 nM on average. This concentration steadily increased to over 500 nM for the largest cells, which are close to the size threshold for mitotic entry. Thus, wild type cells enter mitosis when Cdc25 has accumulated to slightly over 500 nM in the nucleus (Figure 4C).

As a second extension, we measured Cdc25-mNG concentration in *cdr2*Δ cells, which are larger than wild type cells. Consistent with Cdc25 acting as a “sizer” molecule, we measured the same nuclear Cdc25-mNG concentration for wild type and *cdr2*Δ mutants within a given size bin, and *cdr2*Δ cells reached a higher level of Cdc25 in the nucleus because they divided at a larger size (Figures 4D and 4E). Finally, we asked what happens to Cdc25 nuclear concentration when cell growth is inhibited. Treatment of cells with Latrunculin-B (LatB) causes disassembly of the actin cytoskeleton and abolishes cell expansion (Spector et al., 1983; Pan et al., 2014). We found that Cdc25-mNG nuclear accumulation plateaued upon treatment of cells with LatB (Figures S3G and S3H). Put together, these results show that Cdc25 accumulates in the nucleus as a volume-dependent sizer molecule. This scaling property is distinct from Cdr2, which accumulates in the cell middle with cell surface area, indicating that cells have at least 2 distinct pathways monitoring different aspects of size: cell volume (Cdc25) and cell surface area (Cdr2).

### Cdc13 accumulates in the nucleus as a molecular timer

Next, we performed a similar set of experiments for Cdc13-mNG expressed from the endogenous promoter and chromosomal locus. This strain does not exhibit any defects previously suggested for C-terminally tagged constructs, and we confirmed key results with an internally tagged Cdc13-sfGFPint strain (Figures S4A, S4B, and S4G) (Kamenz et al., 2015). We imaged Cdc13-mNG and BFP-NLS in wild type, *rga4Δ*, and *rga2*Δ strain background and plotted Cdc13-mNG intensity as a function of cell surface area or volume. Cdc13-mNG nuclear intensity scaled more closely with cell surface area than with cell volume, indicating a distinct accumulation mechanism from Cdc25 (Figures 5A and S4C-E). The molar concentration of Cdc13 in the nucleus increased from approximately 1 μM in small cells to 3 μM in larger cells (Figure 5B). Surprisingly, Cdc13-mNG did not show properties of a “sizer” molecule when we compared wild type and *cdr2*Δ cells, which are larger than wild type. Instead, *cdr2*Δ mutant and wild type cells had similar Cdc13 nuclear concentrations at birth despite their size differences, while cells in the same size range had different Cdc13 nuclear concentrations (Figure 5C). This conclusion was strengthened by the significant difference in Cdc13-mNG nuclear concentration for wild type versus *cdr2*Δ for cells binned by size (Figure 5D). These results are inconsistent with Cdc13 acting as a sizer molecule.

**Figure 5.**
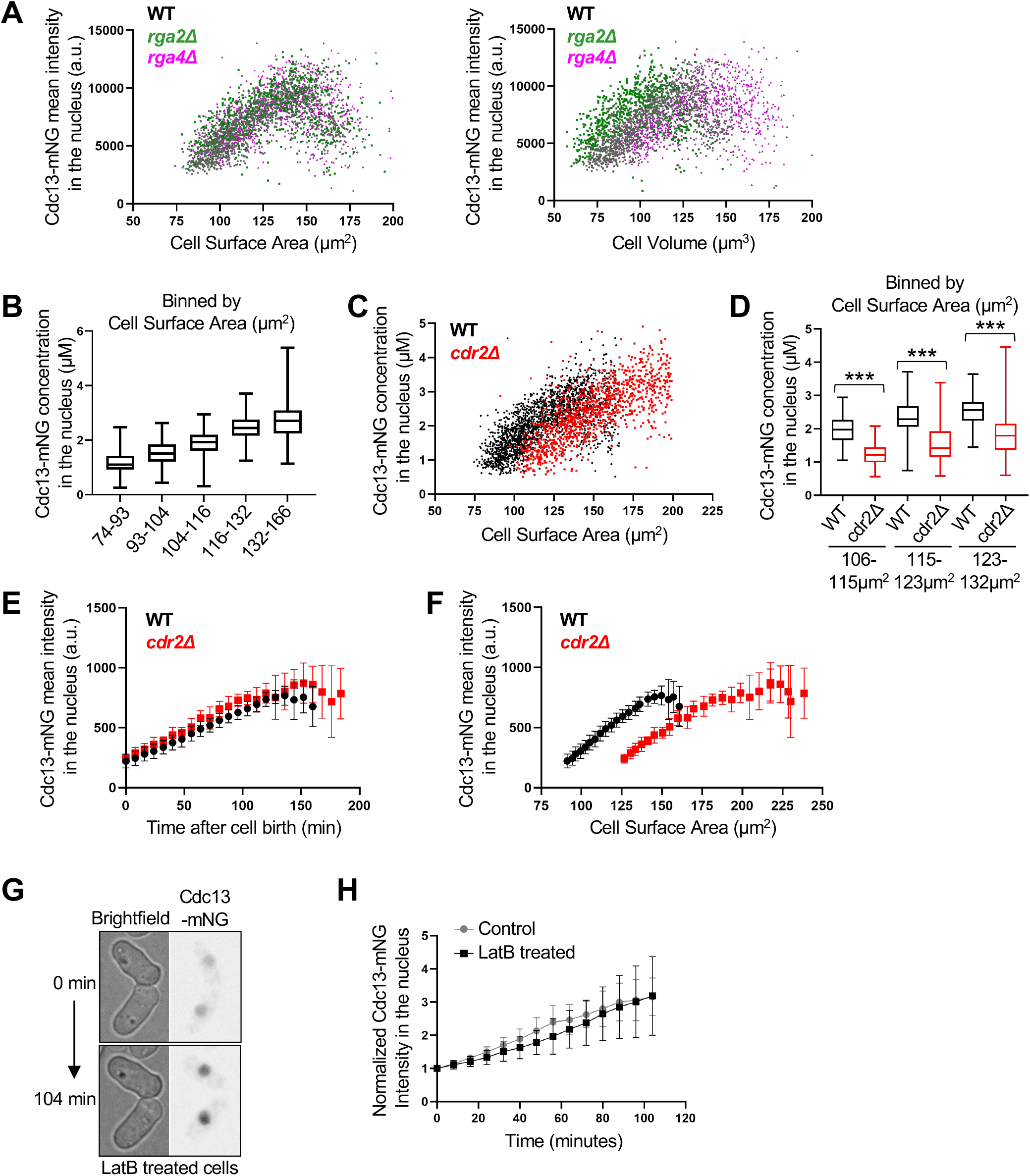
Time-based accumulation of mitotic cyclin Cdc13 in the nucleus. (A) Cdc13-mNG nuclear mean intensity in WT, *rga2*Δ, and *rga4*Δ single cells plotted either by cell surface area or by cell volume (WT n=1,166, *rga2*Δ n=687, and *rga4*Δ n=1,019). The same cells are plotted in the two graphs. (B) Cdc13-mNG nuclear concentration (μM) in WT cells binned by cell surface area. n>280 for each size bin. Graph shows median as a line, quartiles, max, and min. (C) Cdc13-mNG nuclear concentration (μM) in WT and *cdr2*Δ cells plotted by cell surface area (WT n=1,950; *cdr2*Δ n=1,141). (D) Cdc13-mNG concentration (μM) in the nucleus of WT and *cdr2*Δ cells binned by cell surface area (n>75 for each size bin). ***p<0.0001. Graph shows median as a line, quartiles, max, and min. (E-F) Average Cdc13-mNG mean intensity in the nucleus of single cells from time-lapse imaging WT or *cdr2*Δ cells. n=10 cells per strain. Error bars indicate SD. Cdc13-mNG nuclear concentration was plotted either by time since cell birth (panel E) or by cell surface area (panel F). (G) Representative time-lapse image of Cdc13-mNG cells upon addition of 100μM Latrunculin B, an actin inhibitor that halts cell growth. (H) Mean nuclear Cdc13-mNG intensity over time from timelapse imaging of cells treated with or without 100μM Latrunculin B. n=10 cells per experiment. Error bars indicate SD.

We considered the alternative hypothesis that Cdc13 accumulates as a “timer” molecule, such that its concentration reflects time since cell birth as opposed to absolute cell size. Because cells increase in size over time during normal growth conditions, this mechanism could explain accumulation of Cdc13 in our initial experiments as well as in previous studies (Patterson et al., 2019; Curran et al., 2022). To test this idea, we performed timelapse microscopy of Cdc13-mNG in wild type and *cdr2*Δ mutant cells (Figure S5). For each cell, we plotted Cdc13-mNG nuclear concentration as a function of either cell size or time since cell birth. We observed striking overlap of Cdc13 nuclear concentration in wild type and *cdr2*Δ cells when plotted as a function of time since cell birth (Figure 5E). In contrast, there was no overlap between these strains when plotted by cell size (Figure 5F), consistent with our static imaging experiment (Figure 5C and 5D). These combined results are consistent with a model where newborn cells accumulate Cdc13 over time, regardless of their size.

As a final test of Cdc13 acting as a timer molecule, we stopped cell growth by disrupting the actin cytoskeleton with LatB. If Cdc13 acts as a timer molecule, then we expect Cdc13 nuclear concentration to continue increasing in the absence of cell growth. Alternatively, if Cdc13 acts as a sizer molecule like Cdc25, then its nuclear concentration should plateau in the absence of growth.

We found that the nuclear concentration of Cdc13-mNG (and Cdc13-sfGFPint) continued to increase after LatB addition. The rate of this increase was similar to control-treated cells (Figures 5G-H and S4G). Thus, the nuclear accumulation of Cdc13 is time-dependent but not size-dependent. We conclude that Cdc13 is a molecular “timer.”

### The integrated cell size control network can exhibit sizer and adder behaviors

Cell size homeostasis can be revealed by plotting the birth size of single cells against their size increase during the cell cycle from birth to division (Figure 6A). In these plots, a slope of −1 indicates a sizer mechanism, where cells divide at a fixed target size. A slope of 0 indicates an adder strategy, where cells add a fixed amount of size each division. A slope of+1 suggests timer behavior, where cells grow for a fixed duration of time. Previous work established that fission yeast cells exhibit strong sizer behavior in such experiments with a slope near −1 (Fantes, 1977; Sveiczer et al., 1996). Our work combined with previous studies shows that the fission yeast cell size control system is comprised of at least 3 distinct input signals: cell surface area (Cdr2), cell volume (Cdc25), and time (Cdc13). We formalized this integrated system in a qualitative mathematical model that includes size-dependent inhibition of Wee1 (i.e. Cdr2 pathway), size-dependent accumulation of Cdc25, and time-dependent accumulation of Cdc13 (Figure 6B, see Methods for additional details). We found that a three-component model derived from two size-dependent inputs and one time-dependent input is sufficient to recapitulate fission yeast cell size homeostasis with a slope of −1 (Figure 6C). Next, we changed the model to predict the behavior of cells upon loss of a size-dependent input. We altered Wee1 activity from a size-dependent parameter to a constant, size-independent value. This two-component model, consisting of one size-dependent input and one time-dependent input, qualitatively predicted adder-like behavior based on size-dependent Cdc25 activation and time-dependent Cdc13 activation (Figure 6C). Thus, changing from a three-component model to a two-component model can shift size homeostasis from sizer to adder behavior, at least in principle.

**Figure 6.**
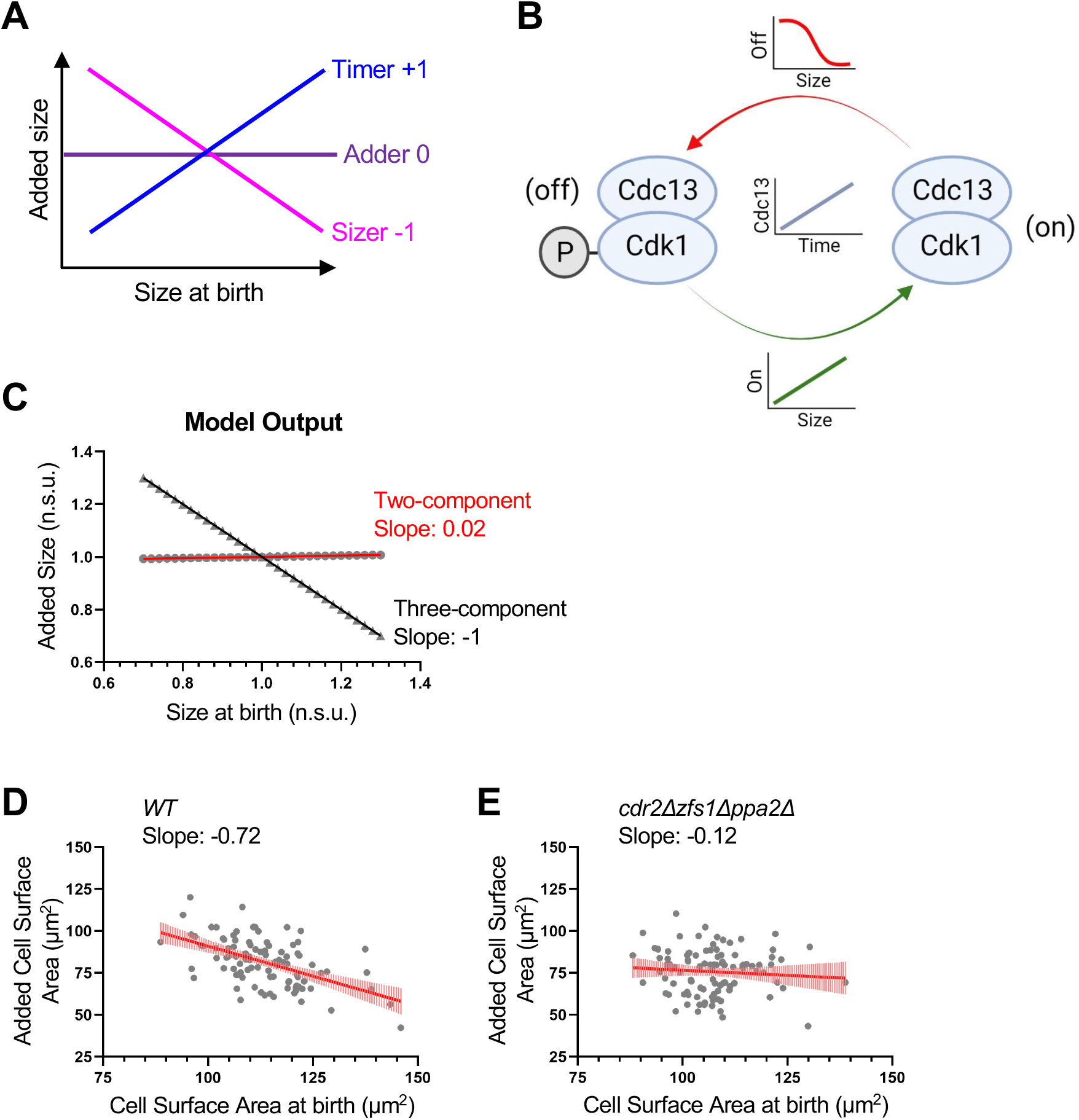
Modeling and experiments identify conditions for adder-like cell size homeostasis. (A) Cell size homeostasis graphic showing the expected slope obtained from plotting the birth size of single cells using timer, adder, or sizer strategies against their size increase during the cell cycle. (B) Schematic of three-component model with time-dependent accumulation of Cdc13 and size-dependent activation and inhibition of Cdk1-Cdc13 complex. (C) Size homeostasis plot from model. A three-component model with 2 sizers and 1 timer reveals sizer behavior (Slope of −1). A two-component model with 1 sizer and 1 timer leads to adder behavior (Slope of 0.02). Added size and size at birth are normalized to the mean added size or mean size at birth, respectively. Normalized size units (n.s.u.). Slope of linear regression displayed. (D-E) Size homeostasis plots of WT (D) and *cdr2*Δ *zfs1*Δ *ppa2*Δ (E) cells. Slope of linear regression line (red) displayed below cell type for each graph. Plots show cell surface area at birth and added surface area for individual cells from timelapse imaging. n>100 for each cell type.

To test this modeling prediction experimentally, we generated cell size homeostasis plots from timelapse microscopy. We measured cell surface area at birth and surface area addition per cell cycle for each cell in a timelapse experiment. Wild type cells exhibited size homeostasis as shown by a slope of −0.72 (Figure 6D), which confirms previous results based on cell length (Fantes, 1977; Scotchman et al., 2021). To remove size-dependent inhibition of Wee1, we used *cdr2*Δ *zfs1*Δ *ppa2*Δ cells. This strain removes the size-dependent Cdr2 pathway and maintains cells at a similar overall size to wild type (Figure 2B). Remarkably, *cdr2*Δ *zfs1*Δ *ppa2*Δ cells exhibited a slope of −0.12 in cell homeostasis plots, indicating near-adder behavior (Figure 6E). We note that both *cdr2*Δ and *zfs1*Δ *ppa2*Δ strains had weakened size homeostasis on their own (slopes of −0.5 and −0.6, respectively) (Figure S6). We conclude that cell size homeostasis can occur through the integration of two molecular sizers and one molecular timer in our model. Removing one molecular sizer from can shift the system to adder-like behavior, revealing plasticity in the size control system due to multiple input pathways.

## DISCUSSION

Our work leads to a new, integrated model for cell size control in fission yeast. We propose that the size control network is comprised of at least three different pathways that monitor distinct aspects of cell growth (Figure 7). First, we have shown that the nuclear concentration of Cdc25 accumulates with cell volume. Second, we found that the nuclear concentration of cyclin Cdc13 increases over time, as opposed to size. Third, previous work has established that Cdr2-Wee1 signaling relates to cell surface area through the spatial density of Cdr2 nodes at the medial cell cortex (Pan et al., 2014; Facchetti et al., 2019). Such an integrated model likely explains why past studies have failed to identify a single “sizer” molecule or pathway in fission yeast. Each of these pathways has been shown to change activity with cell size and growth, but loss of a single pathway does not appear to abolish size homeostasis. Our results do not exclude the possibility of additional pathways and components that monitor other aspects of cell size and growth. For example, we found that reducing the size of *cdr2*Δ mutant cells restored surface area-based divisions, which suggests that additional pathways act with Cdr2 to monitor cell surface area. The pathways that we studied all converge on Cdk1, with both Cdr2-Wee1 and Cdc25 focused on Cdk1-pTyr15. Many mutants alter cell size independently of Cdk1-pTyr15 and could represent more size-dependent pathways in this network (Coudreuse & Nurse, 2010; Navarro & Nurse, 2012). Further, phosphatases that oppose Cdk1 such as PP1 and PP2A could have size-dependent regulation in addition to their known inputs from nutritional availability (Chica et al., 2016; Martín et al., 2017).

**Figure 7.**
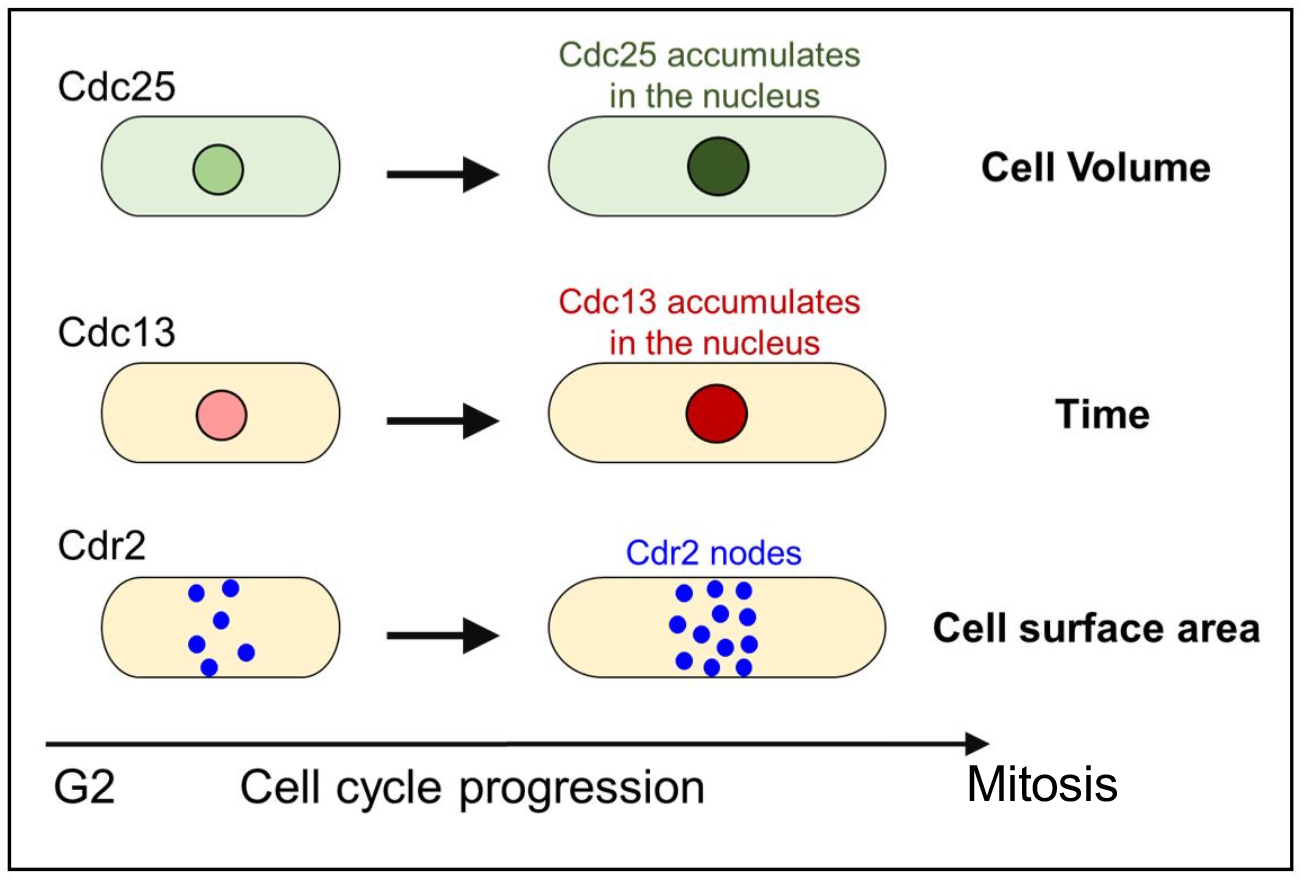
An integrated model for geometry-based cell size control. The fission yeast cell size control network integrates multiple signaling pathways that report on distinct aspects of cell size and growth, including cell volume (Cdc25), time (Cdc13), and cell surface area (Cdr2).

### Size control as an adaptable property

We identified two forms of plasticity within the fission yeast mitotic size control system. First, cells divide at a threshold surface area in one size regime but shift to dividing at a threshold volume at larger sizes. This property was initially identified for *cdr2*Δ mutants (Facchetti et al., 2019), but we have shown that it applies as a general size-dependent behavior. This size-dependent shift is likely to involve changes in the balance or integration of the Cdc25, Cdr2, and Cdc13 pathways. Both Cdc25 concentration and Cdr2 localization continue to increase in large cells (Keifenheim et al., 2017; Sayyad & Pollard, 2022), but a quantitative comparison of their scaling properties could reveal mechanisms for the shift to volume-based division. We also identified a role for ploidy in this shift, consistent with well-known links between cell size and genome content. This finding could relate to ploidy-dependent limits on overall transcription rates in large cells, which also exhibit cytoplasmic dilution and signs of senescence (Zhurinsky et al., 2010; Neurohr et al., 2019; Lanz et al., 2022).

A second form of plasticity in the size control system is the shift from sizer to adder mechanisms for size homeostasis. Size homeostasis plots identify mechanisms that operate across the entire cell cycle and could arise from the combination of multiple pathways. For example, our modeling and experiments identified adder-like behavior upon loss of size-dependent regulation of Wee1, at least for the *cdr2Δ ppa2Δ zfs1*Δ mutant. According to our model, this adder-like behavior can be explained by a molecular sizer (Cdc25) and a molecular timer (Cdc13). This situation appears related to budding yeast, where separate mechanisms for a G1/S sizer and a S/G2/M timer combine to form a phenomenological adder (Chandler-Brown et al., 2017). Both Cdc25 and Cdc13 act on the G2/M transition, meaning that adder-like behavior can arise from the combination of a sizer and a timer that act on the same cell cycle stage.

### Molecular pathways in cell size control

We identified volume-dependent Cdc25 accumulation and time-dependent Cdc13 accumulation as critical pathways in the size control system. Identifying the molecular mechanisms for their accumulation in the whole cell and in the nucleus are important goals for future studies. Previous work has shown that Cdc25 mRNA levels superscale with cell size (Keifenheim et al., 2017; Moreno et al., 1990), raising questions about size-dependent loading of transcription factors and RNA Po1II at the *cdc25+* promoter. *cdc13+* mRNA levels do not oscillate during the cell cycle in large-scale gene expression studies (Rustici et al., 2004), and the *cdc13+* promoter is not required for superscaling of Cdc13 protein (Patterson et al., 2019). Thus, post-transcriptional mechanisms might predominate in Cdc13 superscaling.

Beyond protein levels, both Cdc25 and Cdc13 are more concentrated in the nucleus than in the cytoplasm, which highlights the role of nucleocytoplasmic shuttling in cell size control. Mutations in multiple nuclear shuttling factors lead to changes in cell size (Chua et al., 2002; Moris et al., 2016), but it is currently unknown if shuttling rates are cell size-dependent for Cdc25, Cdc13 or other proteins in this system. The molecular mechanisms that control Cdc25 and Cdc13 accumulation and nuclear concentration likely represent additional layers of the integrated cell size control network.

### Implications for other cell types and organisms

Our work exploited the simple geometry of fission yeast to uncover pathways that monitor distinct aspects of cell geometry and growth. Many other eukaryotic cell types exhibit cell size control, but more complex cell shapes complicate attempts to distinguish between cell volume, cell surface area, and other aspects of size. Despite these technical challenges, it will be interesting to learn if a similar integrated system operates in other eukaryotic cell types and organisms. We note that an integrated system opens possibilities for more complex sizing strategies. For example, bacteria have been shown to use the ratio between surface area and volume as a key parameter for size control (Harris & Theriot, 2016, 2018; Ojkic et al., 2019). Future work on additional signaling pathways that are modulated as cells grow could lead to new insights in this area.

### Limitations of the study

Our experiments were performed under rich nutrient conditions, but both cell size and growth rate are highly dependent on environmental conditions. This raises the possibility that cells might shift the mode of size control under different growth conditions. Further, environmental stress and nutrient limitation are known to impact signaling by the Cdr2, Cdc25, and Cdc13 pathways (Lopez-Girona et al., 1999, 2001; Kelkar & Martin, 2015; Lucena et al., 2017; Allard et al., 2019; Gallardo et al., 2020). It will be interesting to learn how changes to these pathways influence the importance of surface area, volume, and time in the cell size homeostasis network. We also note that we have measured protein concentrations without knowing the added layer of protein activity, such as the catalytic activity of Cdc25 phosphatase. Efforts to measure the specific activity of these proteins from cells of different sizes will extend our understanding of the integrated cell size control system.

## STAR METHODS

### Strain construction and growth conditions

Standard methods were used to grow and culture *Schizosaccharomyces pombe* cells (Moreno et al., 1991). Yeast strains used in this study are listed in Supplemental Table S1. All gene fusions were expressed from their endogenous promoters at the endogenous chromosomal loci. One-step PCR-based homologous recombination was used for C-terminal tagging or deletion of genes on the chromosome (Bähler et al., 1998).

### Microscopy

#### Analysis of cell geometry and fluorescent protein intensity from static images

Fission yeast cells were grown at 25°C in YE4S medium to logarithmic phase for imaging as previously described (Opalko et al., 2022). Cells were placed in a coverglass-bottom dish (P35G-1.5-20C; MatTek), and covered with a piece of YE4S agar prewarmed to 25°C. Imaging was performed using a spinning disk confocal microscope: Yokogawa CSU-WI (Nikon Software) equipped with a 60× 1.4-NA CFI60 Apochromat Lambda S objective lens (Nikon); 405-, 488-, and 561-nm laser lines; and a photometrics Prime BSI camera on an inverted microscope (Eclipse Ti2; Nikon). Multiple fields of view per cell type were imaged within 1 h at room temperature. Images were captured with 27 z-stacks and 0.2-μm step size.

#### Microscopy for cell geometry and intensity measurements over time

For imaging Cdc13 or Cdc25 accumulation over time, cells were mounted on a coverglass-bottom dish (P35G-1.5-20C; MatTek) and covered with a piece of YE4S agar prewarmed to 25°C. Images were acquired with 7 z-stacks and 0.5-μm steps every 8 minutes using a spinning disk confocal microscope (described above). For time-lapse imaging Latrunculin B treated cells, 100μM LatB was added to cells prior to their mounting on a coverglass-bottom dish. YE4S agar containing 100μM LatB was used to cover the cells.

#### Microscopy for cell size homeostasis plots

Cells were imaged at room temperature in microfluidic flow chambers (Millipore, CellAsic ONIX, 3.5 - 5.5 μm Y04C-02). First, flow chambers were primed by flowing YE4S media for 7 min at 6 psi. Diluted cells were then loaded into chambers and YE4S media was flowed at 6 psi for the duration of the experiment. Cells were allowed to acclimate in the chambers for 1 hour prior to imaging. Brightfield images of cells were acquired with 3 z-stacks and 0.5-μm steps every 8 minutes using a spinning disk confocal microscope (described above). Cell size at birth/division and cell cycle time were analyzed for 2-3 generations of cells. Cell size at division and cell cycle time were consistent for the duration of this imaging period.

### Cell and nuclei segmentation

Brightfield images were processed for cell segmentation and cell size analysis using a semiautomated pipeline. First using ImageJ, a smoothing filter and Gaussian blur was applied to each optical section of an image to reduce image noise. Next, global thresholding was performed by selecting a grey value maximum to produce a binary image for each z-section. An optical section outside of the middle focal plane with intact boundary bands around cells was selected for further processing. The flood fill tool was used (by hand) to generate a black background so that image pixel values in the background were set to 0 and cells in white were 1. Binary images were further processed by morphological erosion and subsequent dilation operators to remove regions between cell clumps for better single-cell segmentation. Images were also processed to remove cells along the edge of the image. The paintbrush tool was used to further separate clumped cells by hand, and the flood fill tool was used to delete any abnormal cells or unresolved clumps of cells. The resulting binary image (“cell mask”) was compared with the original bright-field image and confirmed to be an accurate representation of cell size.

BFP-NLS images were processed for nuclei segmentation and nuclei size analysis using a semiautomated pipeline. First using ImageJ, a sum projection was created from image z-sections. To reduce image noise, a smoothing filter and Gaussian blur was applied. Next, global thresholding was performed by selecting grey values for the BFP-NLS signal to produce a binary image of nuclei. Images were also processed to remove nuclei along the edge of the image. The resulting binary image (“nuclei mask”) was compared to the original BFP-NLS image and confirmed to be an accurate representation of nuclear size.

To obtain cell masks of dividing cells, morphological erosion of cell and nuclei masks was initially performed using ImageJ. This is done so that nuclei are not touching the cell border and to better separate clumped cells. To obtain a single image with nuclei overlaid on cells, eroded cell and nuclei masks are processed by a MATLAB code that sets image pixel values in the background and nuclei to 0 and cells in white to 1. Nuclei-overlaid cell images are further processed with a MATLAB code that removes interphase cells and creates a binary mask containing only dividing cells (cells with two nuclei, indicating active division). The paintbrush tool was used to edit clumped cells by hand and the flood fill tool was used to delete any abnormal cells or unresolved clumps of cells. Once binary masks were confirmed to contain only dividing cells, by comparison to original BFP-NLS and brightfield images, images were further processed to remove nuclei so the image pixel values for the whole cell is 1 and processed by morphological dilation to return cells to normal size. This resulting binary image is the mask of dividing cells.

For time-lapse images, individual cropped cells are processed like static images with some modifications. To generate cell masks from brightfield time-lapse images, a single z-section outside of the middle focal plane with intact boundary bands around cells was selected for each time-point for further processing. A smoothing filter and Gaussian blur were applied to each image time point to reduce image noise. A sharpening filter was applied to increase contrast of the boundary band around cells, followed by a Sobel edge detector to highlight the sharp change in intensity of the boundary band. Next, global thresholding was performed by selecting a grey value that includes the boundary around cells to produce a binary image for each time point. The flood fill tool was used (by hand) to generate a black background so that image pixel values in the background were set to 0 and cells in white were 1. The paintbrush tool was used to further separate clumped cells by hand or edit cell boundary. The resulting cell mask of each time point was compared with the original brightfield image and confirmed to be an accurate representation of cell size. To create nuclei masks from the BFP-NLS channel for each time point, the same semiautomated pipeline was used to generate nuclei masks (above). Each step in the pipeline was applied to all BFP-NLS image time points.

### Cell geometry measurements

For each strain, the cell width was manually measured from the cell mask in ImageJ using the straight-line tool. The average cell width divided by 2 (cell radius) was determined for a population of cells (n > 100). We assumed that each cell in a given strain had the same cell radius (average of population) for calculation of cell surface area and volume (Table 1). In MATLAB, the cell length or cell symmetry axis of individual segmented cells (from cell masks) was identified by principal component analysis of the cloud points internal to the cell (Facchetti et al., 2019). Cell surface area or volume of individual cells was calculated in MATLAB using the equation for surface area and volume of a cylinder with hemispherical ends, because of the rod-like shape of fission yeast cells. In MATLAB, the major and minor axis of nuclei (from nuclei mask) was obtained using the *regionprops* function to return the length (in pixels). To get the actual length, we multiplied the number of pixels by the length in microns represented by one pixel. Volume of nuclei was calculated using the equation for an ellipsoid, assuming two of the three semi-axes are equal in length.

**Table 1.**

|  |  |
| --- | --- |
| Calculation of cell or nuclear surface area and volume |  |
| Cell radius<sup>a</sup> | $R$ |
| Cell length | $L$ |
| Cell surface area | $2\pi RL$ |
| Cell volume | $\pi R^2 L - 2/3\pi R^3$ |
| Nuclei minor axis | $mj$ |
| Nuclei major axis | $mn$ |
| Nuclei volume | $1/6\pi * mj * mn^2$ |

<sup>a</sup>Average radius of cell population determined by manual measurements of individual cells. We assume that all cells of a given type have the same cell radius.

For cell size homeostasis studies, brightfield images were analyzed by hand. In ImageJ, the straight-line tool was used to measure the length and width of each cell at birth and division (determined by the presence of a septa). The cell cycle time from birth to division for each cell was denoted. Cell surface area of individual cells was calculated using the equation for a cylinder with hemispherical ends.

### Measurement of fluorescent intensity

Summed intensity projections of z-sections were used for analysis of fluorescent intensity. MATLAB codes were used to measure mean intensity or sum intensity of fluorescent proteins in the cytoplasm, nucleus, or whole cell (as defined by cell and nuclei masks). In supplemental graphs (Figure S3B-C; Figure S4D-E), Cdc25-mNG or Cdc13-mNG values are the sum intensity in the nucleus divided by the nuclear volume for single cells.

For analysis of Cdc13-mNG, Cdc13-sfGFPint, and Cdc25-mNG intensity in the nucleus after treatment with LatB (Figure 5H; Figure S3H; Figure S4G), the lowest intensity value was normalized to 1 and plotted on graphs.

### Calculation of protein concentration (μM) from fluorescent intensities

Molecule counting was performed as described in Wu and Pollard (2005). A standard curve relating fluorescent intensity to concentration (μM) was created by imaging mNG-tagged Fim1, Arp2, Spn1, Mid1, and Cdc12, which have known concentrations. We imaged Cdc13-mNG and Cdc25-mNG in the same genetic background using the same imaging conditions. Cells were grown at 25°C in YE4S medium to logarithmic phase for imaging. Cells were placed on a coverglass-bottom dish and covered with a piece of YE4S agar prewarmed to 25°C. Imaging was performed using a spinning disk confocal microscope (described above). For each strain, images were captured with 27 z-stacks and 0.2-μm steps. Sum projections of all 27 z-stacks were used for analysis. To make the standard curve, mean fluorescent intensity values of mNG-tagged proteins was measured in the whole cell (and nucleus for Cdc13-mG and Cdc25-mNG). Fission yeast cells that do not contain any mNG tagged proteins were also measured and the average intensity value from 100 cells was used for background subtraction. Mean fluorescence values for Fim1, Arp2, Spn1, Mid1, and Cdc12 were then plotted against their known whole cell concentrations (μM) to generate the standard curve. Mean fluorescent intensity of Cdc25-mNG or Cdc13-mNG were plotted against the best-fit line of the standard curve to interpolate the nuclear or whole cell concentration (μM).

### Immunoblotting

Cells expressing Cdc13-mNG, Cdc13-mCherry, Cdc13-sfGFPint, or untagged Cdc13 protein were grown in YE4S medium at 25°C or 36.5°C for 5 hours before harvesting. Immunoblot of cell extracts was performed and the presence of Cdc13 was detected using an anti-Cdc13 antibody (Novus Biologicals).

### Mathematical Model

We consider a model where the total intracellular concentration of Cdc13 increases within a cell cycle as per the differential equation

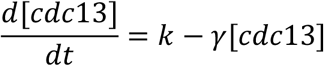

with synthesis rate *k* and dilution rate *γ*. As per this model, the Cdc13 level at a given time *t* within the cell cycle is given by the timer-based equation

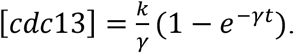

For an exponential increase in cell size *V*

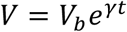

where *V_b_* is the newborn cell size, the total Cdc13 levels can be rewritten as

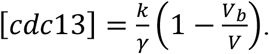

In the *cdr2*Δ background, we assume Cdc13 activation occurs with a rate *k*_1_*V* proportional to size (due to Cdc25 concentration increasing linearly with size), and deactivation occurs at a constant but large rate *k*_2_. Assuming fast cycles of activation/deactivation relative to the time scale of cell growth yields the following concentration of activated Cdc13

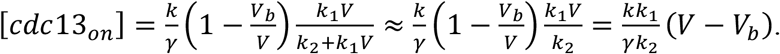

If mitotic entry is driven by [*cdc*13_*on*_] reaching a critical threshold *T*, then it is easy to see from the above equation that it will follow an adder-based size control mechanism

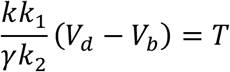

where the size added from cell birth to division *V_d_* – *V_b_* becomes invariant of *V_b_*.

In Fig. 6, the volume added *V_d_* – *V_b_* is plotted as a function of newborn size *V_b_* as it is varied between 0.7 and 1.3, where here 1 unit corresponds to the average newborn size. Given *V_b_*, the cell division size *V_d_* is obtained by solving

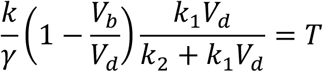

which can be rewritten as

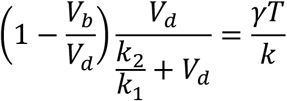

assuming 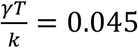 and 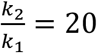. Note with these parameters when *V_b_* = 1 then *V_d_* = 2.

To realize a sizer-based cell size control, we consider a size-dependent deactivation rate *k*_2_ (*V*) that monotonically decreases with increasing *V*. To realize a perfect sizer-based mitotic entry we consider a threshold function that switches from a high to low value when the cell reaches the critical size. More specifically, in Fig. 6 we plot the solution *V_d_* to

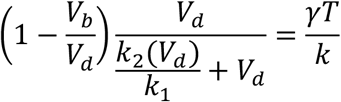

considering 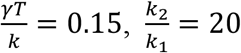 when *V_d_* <2 and 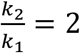 when *V_d_* > 2.

Model code available at https://github.com/millerk89/sizer-timer-pombe

### Quantification and Statistical analysis

For Figure 1G, 4B, 4E, 5D, and S3H Data analysis was performed using Prism 8 (GraphPad Software). A two-tailed Student’s t test was performed to determine statistical differences between two sets of data. An ANOVA was performed to determine statistical differences between sets of data.

### Overlap analysis calculations

We adapted the overlap index from previous work (Pastore and Calcagnì, 2019). We used the overlapping index to compute similarity between cell surface area or volume at division of two yeast strains (data sets). The overlapping index is defined as per

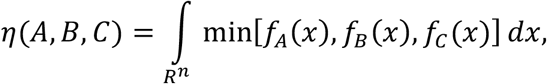

where *f_A_*(*x*) and (*x*) are probability densities of the data sets A and B, respectively. This index is computed using the R library overlapping, available at https://cran.r-project.org/package=overlapping. The index was bootstrapped 1000 times to compute confidence intervals. We extended the overlapping index to compute similarity between three strains as per

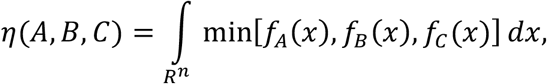

where *f_c_*(*x*) represents the estimated density of surface area/volume for the strain C dataset. This modification is included in our codes as an explicit R function definition that includes the modification of the code available in https://cran.r-project.org/package=overlapping. We performed comparisons between overlapping index of strains for both surface area and volume probability densities, checking surface area or volume overlapping index by bootstrapping, and counting the number of resampling surface area or volume overlapping index is greater. The p values for all overlap index comparisons of surface area and volume were less than 0.001, indicating a significant difference. An exception is Figure 1H where cells are treated with 0.02μM 1-NM-PP1.

### Binning analysis

In Figures 4B, 4E, and 5D we binned the cells by their cell surface area or volume with the same number of cells in each bin using the quantile function included in R.

## Abbreviations

WT: wild type;
mNG: mNeonGreen;
NLS: Nuclear localization sequence;
BFP: blue fluorescent protein;
Latrunculin B: LatB;

## ACKNOWLEDGEMENTS

We thank members of the Moseley laboratory for helpful discussions and comments on the manuscript; as well as the Biomolecular Targeting Core (BioMT) (P20-GM113132) for use of equipment; and Damien Coudreuse, Silke Hauf, and Sophie Martin for sharing yeast strains. This work was funded by grants from the National Institutes of General Medical Sciences (NIGMS) (R01GM099774 and R01GM133856) to J.B.M.

## AUTHOR CONTRIBUTIONS

Conceptualization, K.E.M. and J.B.M.; Methodology, K.E.M., C.V.-G., A.S., and J.B.M.; Software, C.V.-G. and A.S.; Formal Analysis, K.E.M. and C.V.-G.; Investigation, K.E.M.; Writing – Original Draft, K.E.M. and J.B.M.; Writing – Reviewing and Editing, K.E.M., C.V.-G., A.S., and J.B.M.; Supervision, A.S. and J.B.M.; Funding Acquisition, J.B.M.

## DECLARATION OF INTERESTS

The authors declare no competing interests

## SUPPLEMENTAL FIGURE LEGENDS

**Figure S1. Semi-automated image analysis pipeline to measure cell and nuclear size along with fluorescent protein intensities.** (A) Workflow for semi-automated image analysis. This example shows *cdc25-mNG BFP-NLS* cells imaged by spinning disk confocal microscopy. ImageJ is used to generate cell masks from brightfield images and nuclear masks from BFP-NLS images. MATLAB is used to segment cells and nuclei, assigning each nucleus to a cell. Our MATLAB code calculates cell/nuclear size (length, width, surface area, volume) as well as fluorescent intensities in the whole cell, nucleus, and cytoplasm. (B) This pipeline was validated using skinny (*rga2*Δ) and wide (*rga4*Δ) mutants to confirm that cells divide at a constant surface area, as opposed to volume. (C) Overlap index analysis for the data in panel B. SA, surface area. Vol, volume. (D) Our pipeline confirms that *cdr2*Δ cells divide at a constant volume, as opposed to surface area (Facchetti et al., 2019). (E) Overlap index analysis for data in panel D. (F) Cell length at division (μm) for the indicated strains and conditions. All graphs show median as a line, quartiles, max, and min.

**Figure S2. Overlap index graphs.** Graphs show probability of cell surface area or volume at division of indicated cell types and are a visual representation of the overlap index for the following experiments: (A) Dividing WT, *rga2*Δ, and *rga4*Δ cells in Figure S1B-C; (B) Dividing *cdr2*Δ cells in Figure S1D-F; (C) Dividing *wee1-2x* cells in Figure 1D-E; (D) dividing *cdc13-L-cdc2as* cells with indicated treatments in Figure 1G-H; (E) dividing *cdr2*Δ *zfs1*Δ *ppa2*Δ cells in Figure 2C-D; (F) dividing *ppa2*Δ cells in Figure 3B-C; (G) dividing diploid cells in Figure 2F-G.

**Figure S3. Additional analysis of Cdc25 nuclear accumulation.** (A) The same plots of Cdc25-mNG nuclear intensity as Figure 4A, but with mitotic cells removed and linear regression lines for each cell type shown. (B) Cdc25-mNG sum intensity in the nucleus of WT, *rga2*Δ, and *rga4*Δ single cells plotted by cell surface area (μm^2^) or cell volume (μm^3^) (WT n=502, *rga2*Δ n=867, and *rga4*Δ n=684). Note that the same result is obtained by plotting sum intensity as by plotting mean intensity. (C) The same plots as Figure S3B but with mitotic cells removed and linear regression lines for each cell type shown. (D) Overlap index analysis for results in main Figure 4B, which show Cdc25-mNG nuclear intensity for cells binned by their cell surface area or volume. SA, surface area. Vol, volume. (E) Standard curve for calculating Cdc13 and Cdc25 concentrations (μM). Error bars indicate SD. Linear regression used to interpolate experimental values. Fim1 and Arp2 were used to generate the standard curve, but are not pictured here. Whole cell concentrations for standard proteins taken from Wu and Pollard (2005). (F) The whole cell concentration (μM) of Cdc25-mNG increases with cell size. Cells are binned by cell surface area (n>250 for each size bin). (G) Representative time-lapse image of *cdc25-mNG* cells grown in the presence of 100μM Latrunculin B (LatB). Cdc25-mNG does not accumulate in the nucleus when growth is inhibited by addition of 100μM LatB, an actin inhibitor. (H) Mean normalized nuclear Cdc25-mNG intensity over time in individual cells treated with or without 100μM LatB. n=10 cells per experiment. Error bars indicate SD. Asterisks indicate significant difference in fluorescent intensity (p>0.05) at given time point.

**Figure S4. Additional analysis of Cdc13.** (A) The indicated strains were spotted with 10× serial dilutions onto YE4S plates. Plates were incubated for 2-4 days prior to imaging. Yeast cells expressing tagged Cdc13 or Cdc25 do not exhibit growth defects. (B) Western blot of whole cell extracts from the indicated strains. Cells were grown at 25°C or 36.5°C for 5 hours before harvesting. Anti-Cdc13 antibody was used to test if fusion proteins were cleaved under experimental conditions used in this study. No untagged or cleaved Cdc13 protein was detected in cells expressing Cdc13-mNG, Cdc13-sfGFP, or Cdc13-mCherry. (C) The same plots of Cdc13-mNG nuclear accumulation as Figure 5A, but with mitotic cells removed and linear regression lines for each cell type shown. (D) Cdc13-mNG sum intensity in the nucleus of WT, *rga2*Δ, and *rga4*Δ single cells plotted by cell surface area (μm^2^) or cell volume (μm^3^) (WT n=1,166, *rga2*Δ n=687, and *rga4*Δ n=1,019). (E) The same plots as Figure S4D but with mitotic cells removed and linear regression lines for each cell type shown. (F) The whole cell concentration (μM) of Cdc13-mNG increases with cell size. Cells are binned by cell surface area. n>280 for each size bin. Graph shows median as a line, quartiles, max, and min. (G) Mean normalized nuclear Cdc13-sfGFPint intensity over time in individual cells treated with or without 100μM LatB. n=10 cells per experiment. Error bars indicate SD.

**Figure S5. Time-lapse imaging of Cdc13-mNG in wild type and *cdr2*Δ cells.** Time-lapse images of Cdc13-mNG, BFP-NLS, and cell dimensions by brightfield in WT (A) and *cdr2*Δ cells (B). Images captured every 8 minutes. Images are inverted maximum projection images, except brightfield images are single middle z sections.

**Figure S6. Additional cell size homeostasis analysis.** Size homeostasis plots of *cdr2*Δ (A), and *zfs1Δ ppa2*Δ (B) cells. Slope of linear regression line (red) displayed below cell type for each graph. n>100 for each cell type.

## Notes

### Competing Interest Statement

The authors have declared no competing interest.

