## Supplemental Figures S1-S6 for "The fission yeast cell size control system integrates pathways measuring cell surface area, volume, and time"

### Figure S1

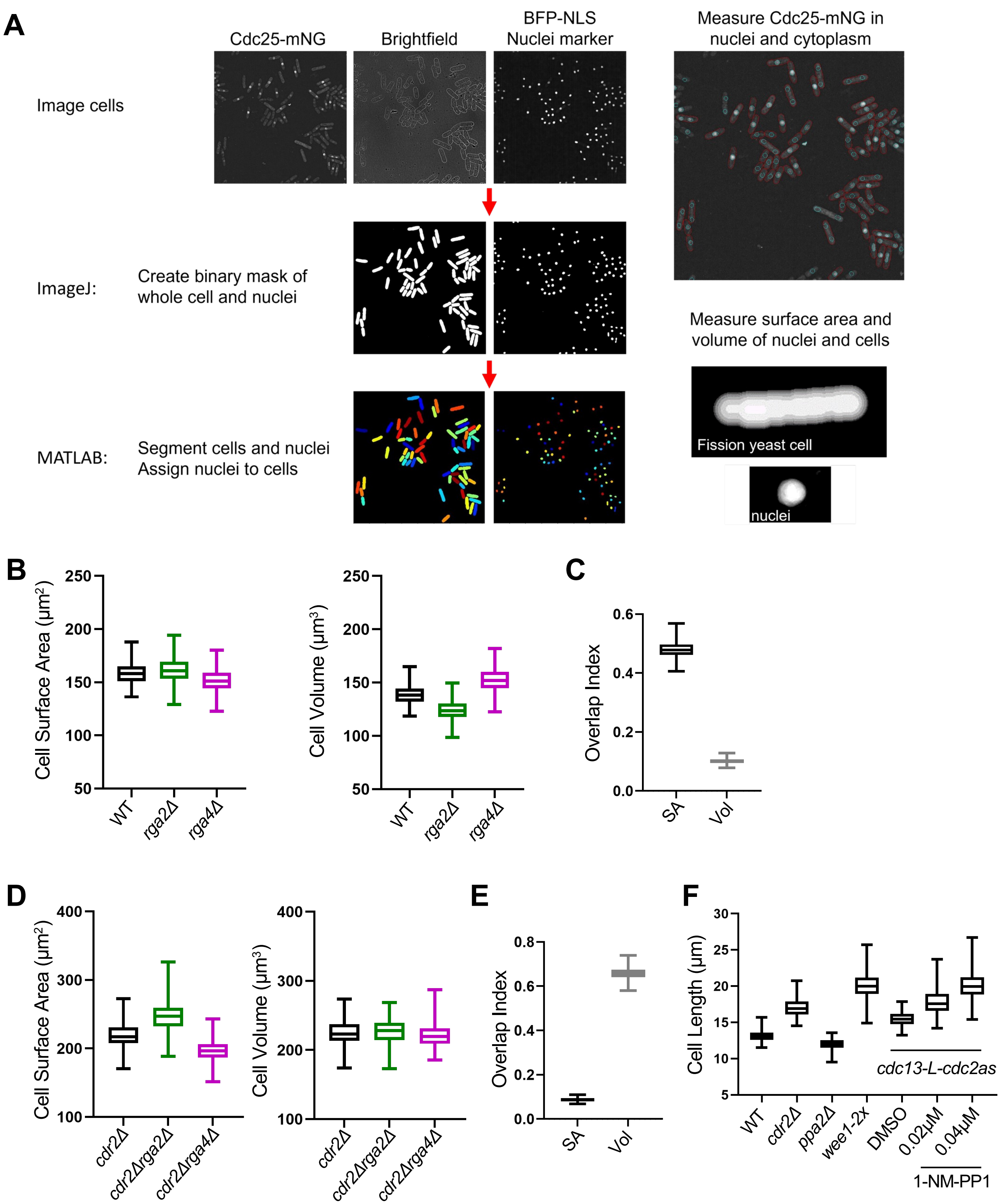
A
Cdc25-mNG
Brightfield
BFP-NLS
Nuclei marker
Measure Cdc25-mNG in
nuclei and cytoplasm
Image cells
ImageJ:
Create binary mask of
whole cell and nuclei
Measure surface area and
volume of nuclei and cells
Fission yeast cell
MATLAB:
Segment cells and nuclei
Assign nuclei to cells
nuclei
B
Cell Surface Area (μm²)
250
200
150
100
50
WT
rga2Δ
rga4Δ
Cell Volume (μm³)
C
Overlap Index
0.6
0.4
0.2
0.0
SA
Vol
D
400
300
200
100
cdr2Δ
cdr2Δrga2Δ
cdr2Δrga4Δ
E
0.8
F
Cell Length (μm)
30
25
20
15
10
5
ppa2Δ
wee1-2x
DMSO
0.02μM
0.04μM
cdc13-L-cdc2as
1-NM-PP1

### Figure S2 – overlap figures

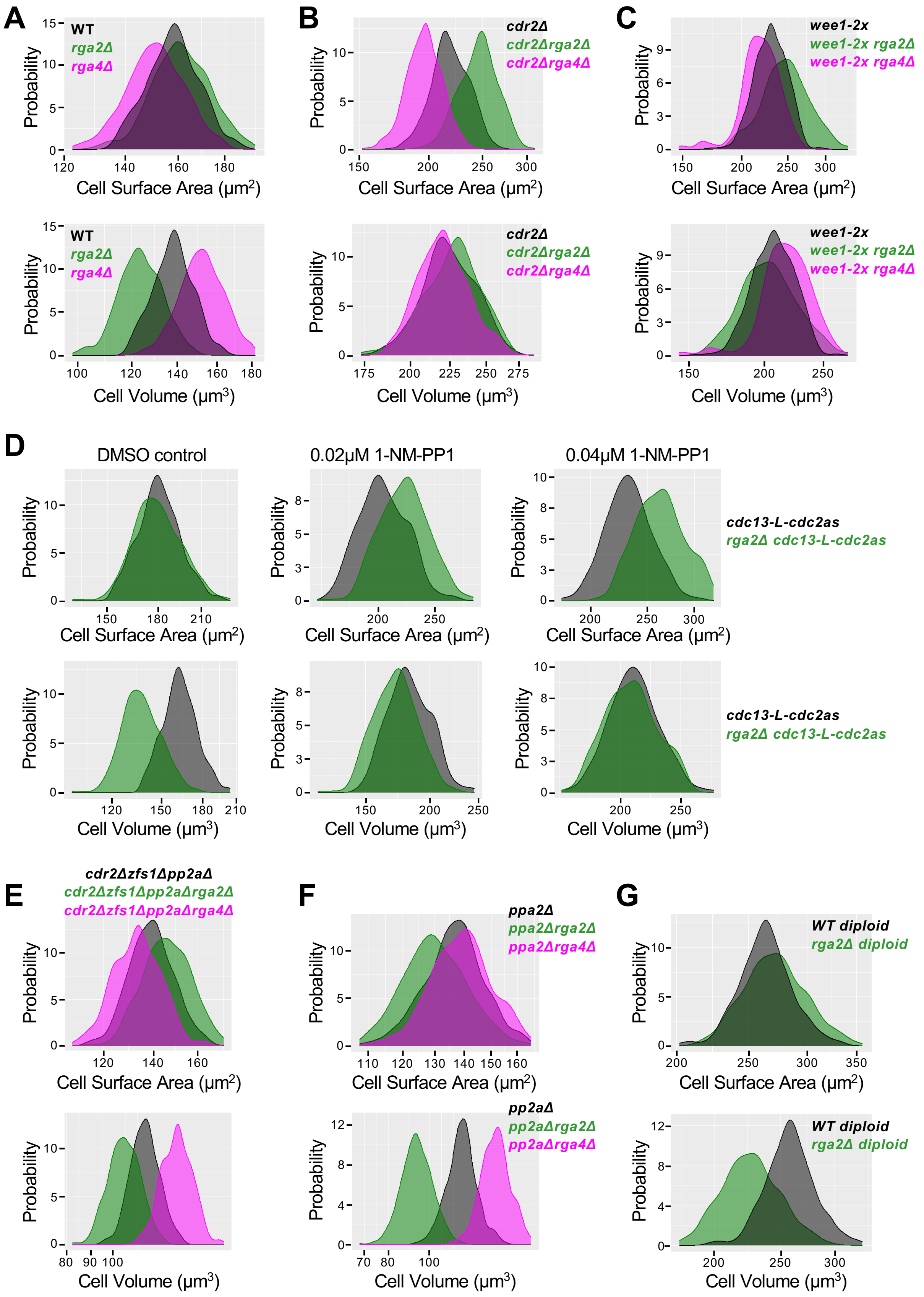

### Figure S3

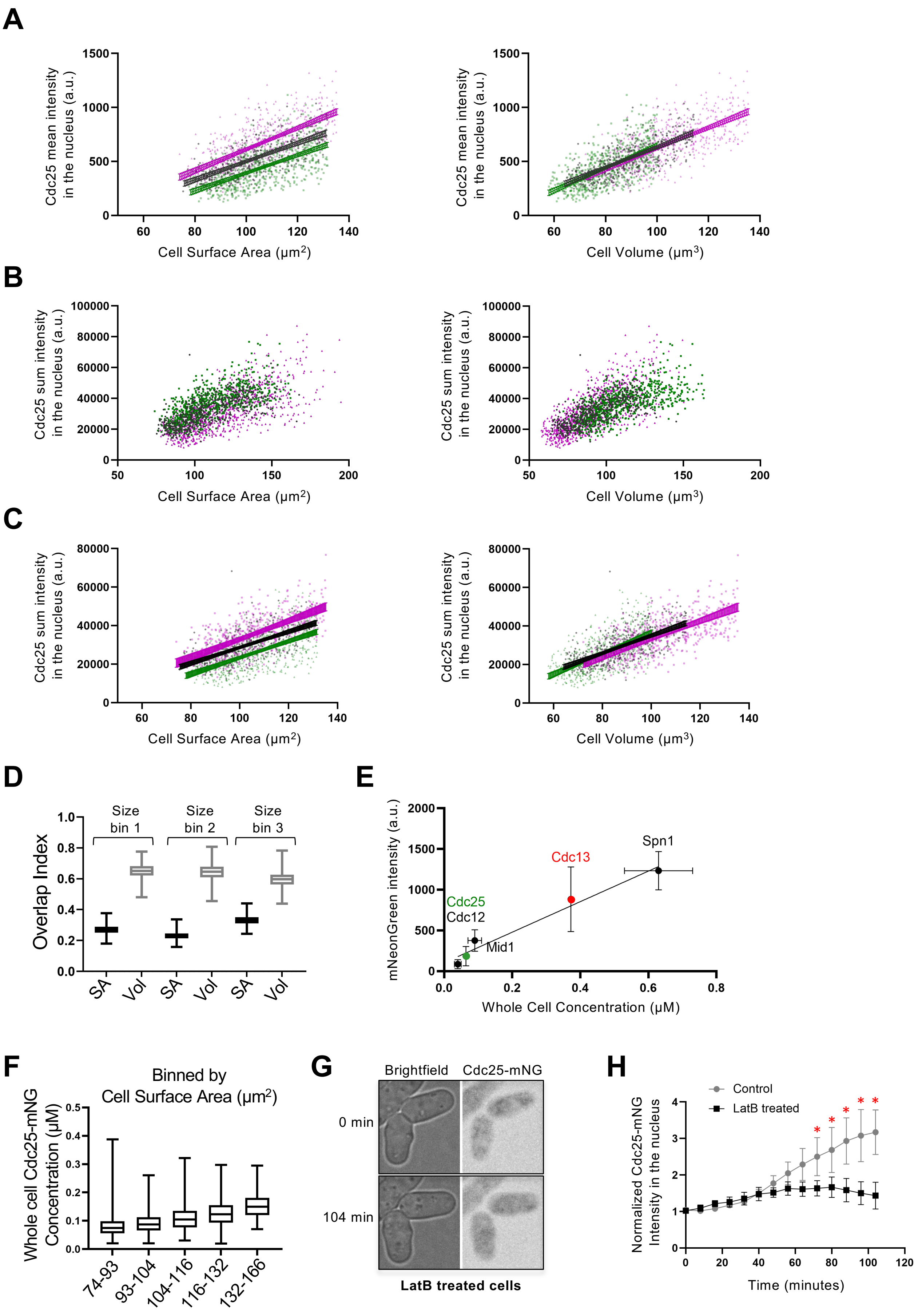
A
Cdc25 mean intensity in the nucleus (a.u.)
Cell Surface Area (μm2)
Cell Volume (μm3)
B
Cdc25 sum intensity in the nucleus (a.u.)
Cell Surface Area (μm2)
Cell Volume (μm3)
C
Cdc25 sum intensity in the nucleus (a.u.)
Cell Surface Area (μm2)
Cell Volume (μm3)
D
Overlap Index
Size bin 1
Size bin 2
Size bin 3
SA
Vol
E
mNeonGreen intensity (a.u.)
Whole Cell Concentration (μM)
Spn1
Cdc13
Cdc25
Cdc12
Mid1
F
Binned by Cell Surface Area (μm2)
Whole cell Cdc25-mNG Concentration (μM)
74-93
93-104
104-116
116-132
132-166
G
Brightfield
Cdc25-mNG
0 min
104 min
LatB treated cells
H
Control
LatB treated
Normalized Cdc25-mNG Intensity in the nucleus
Time (minutes)

### Figure S4

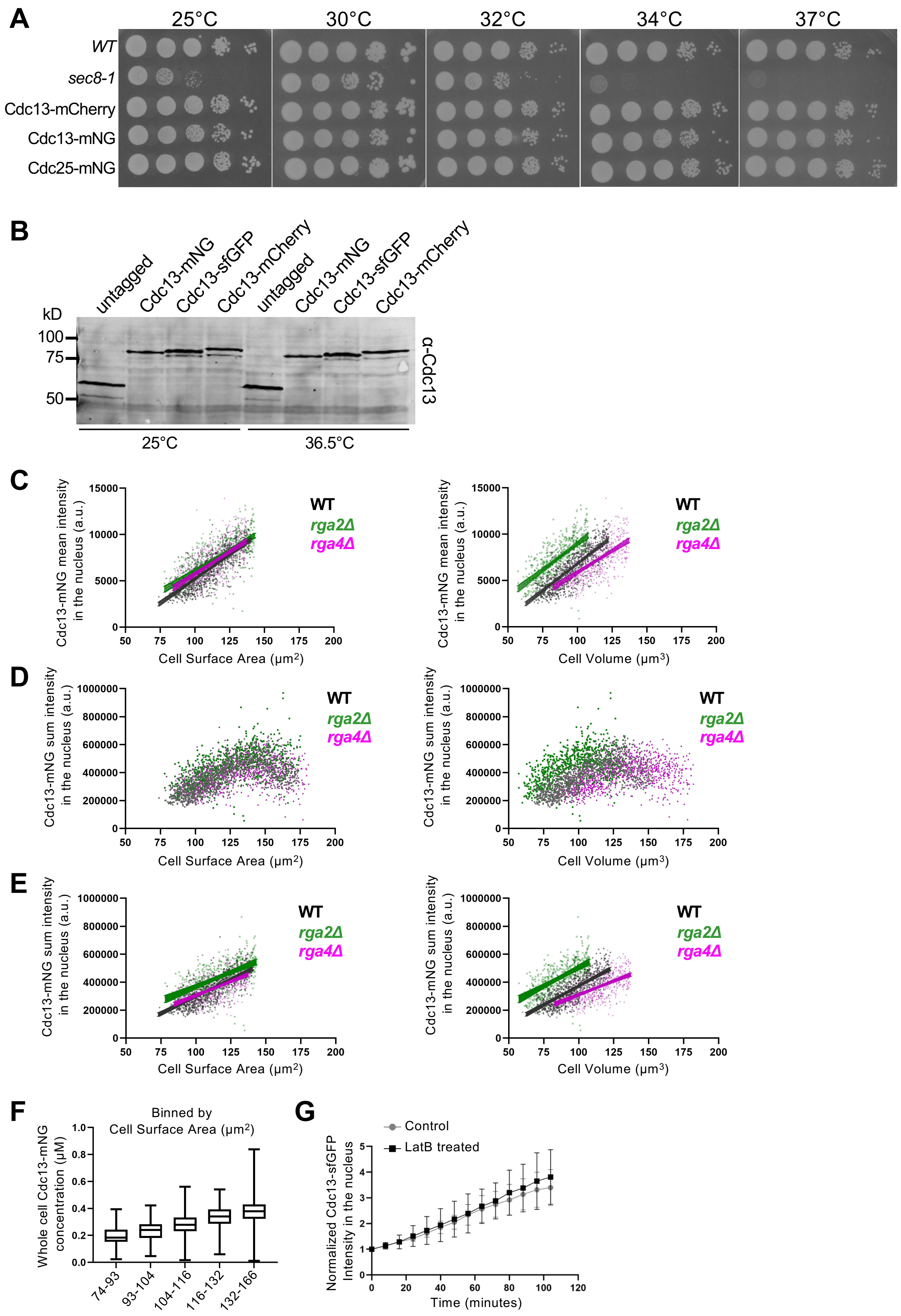

A
25°C
30°C
32°C
34°C
37°C
WT
sec8-1
Cdc13-mCherry
Cdc13-mNG
Cdc25-mNG
B
untagged
Cdc13-mNG
Cdc13-sfGFP
Cdc13-mCherry
kD
100
75
50
α-Cdc13
25°C
36.5°C
C
Cdc13-mNG mean intensity in the nucleus (a.u.)
Cell Surface Area (μm²)
Cell Volume (μm³)
WT
rga2Δ
rga4Δ
D
Cdc13-mNG sum intensity in the nucleus (a.u.)
E
F
Binned by Cell Surface Area (μm²)
Whole cell Cdc13-mNG concentration (μM)
74-93
93-104
104-116
116-132
132-166
G
Control
LatB treated
Normalized Cdc13-sfGFP Intensity in the nucleus
Time (minutes)

### Figure S5

**A**

WT cell

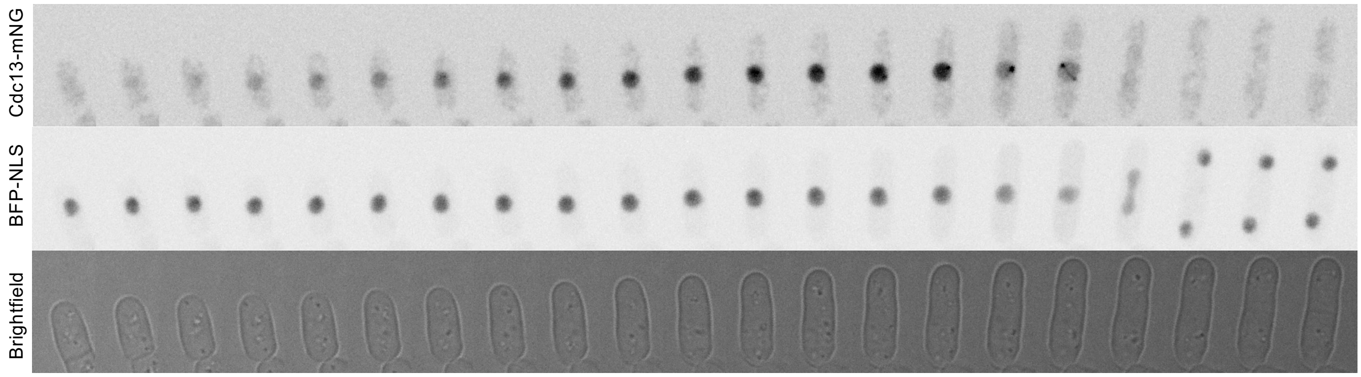


Images captured every 8 minutes

**B**

*cdr2Δ* cell

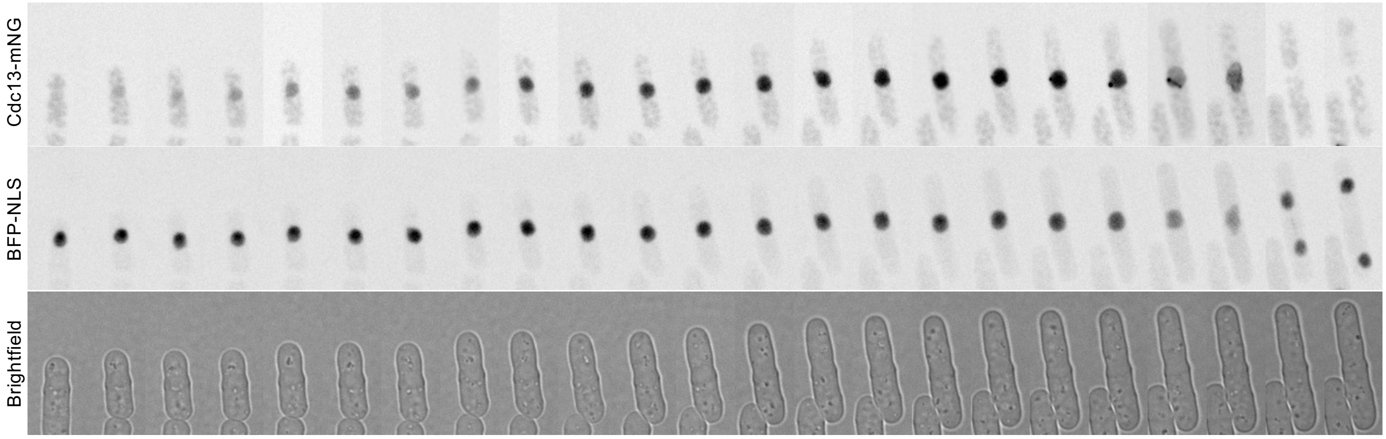


Images captured every 8 minutes

### Figure S6

**A**

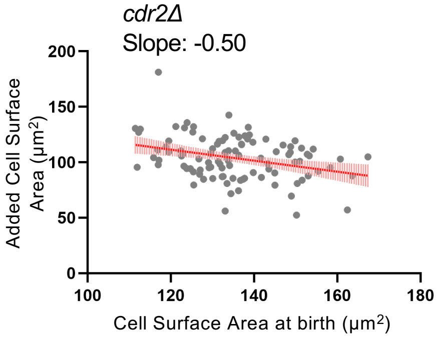

*cdr2Δ*
Slope: -0.50
Added Cell Surface Area (μm2)
0
50
100
150
200
100
120
140
160
180
Cell Surface Area at birth (μm2)


**B**

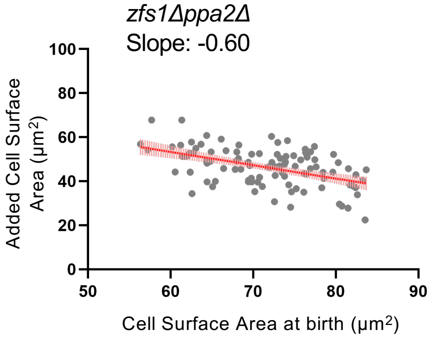

*zfs1Δppa2Δ*
Slope: -0.60
Added Cell Surface Area (μm2)
0
20
40
60
80
100
50
60
70
80
90
Cell Surface Area at birth (μm2)
