## Supplemental Table S1 for "The fission yeast cell size control system integrates pathways measuring cell surface area, volume, and time"

### Table S1. Yeast strains used in this study

| Strain | Genotype | Source |
|---|---|---|
| **JM366** | *972 h- (Alternate name: PN1)* | Lab Collection |
| **JM6972** | *cdc13-yomNeonGreen::hphR lys3+:ptdh1*:NLS-linker-mTagBFP2:terminatortdh1:kanMX leu1:[pJK148-Pact-yomRuby3-term] h+* | This Study |
| **JM7003** | *rga2Δ::kanMX6 lys3+:ptdh1*:NLS-linker-mTagBFP2:terminatortdh1:kanMX cdc13-yomNeonGreen::hphR leu1:[pJK148-Pact-yomRuby3-term] h-* | This Study |
| **JM7008** | *rga4Δ::kanMX6 lys3+:ptdh1*:NLS-linker-mTagBFP2:terminatortdh1:kanMX cdc13-yomNeonGreen::hphR leu1:[pJK148-Pact-yomRuby3-term]* | This Study |
| **JM7088** | *cdr2Δ::NAT lys3+:ptdh1*:NLS-linker-mTagBFP2:terminatortdh1:kanMX cdc13-yomNeonGreen::hphR leu1:[pJK148-Pact-yomRuby3-term] h+* | This Study |
| **JM7171** | *lys3+:ptdh1*:NLS-linker-mTagBFP2:terminatortdh1:kanMX cdc25-yomNeonGreen::hphR leu1:[pJK148-Pact-yomRuby3-term] h-* | This Study |
| **JM7172** | *rga4Δ::kanMX6 lys3+:ptdh1*:NLS-linker-mTagBFP2:terminatortdh1:kanMX cdc25-yomNeonGreen::hphR leu1:[pJK148-Pact-yomRuby3-term] h-* | This Study |
| **JM6752** | *rga2Δ::kanMX6 lys3+:ptdh1*:NLS-linker-mTagBFP2:terminatortdh1:kanMX cdc25-yomNeonGreen::hphR leu1:[pJK148-Pact-yomRuby3-term] h-* | This Study |
| **JM6753** | *cdr2Δ::NAT lys3+:ptdh1*:NLS-linker-mTagBFP2:terminatortdh1:kanMX cdc25-yomNeonGreen::hphR leu1:[pJK148-Pact-yomRuby3-term] h-* | This Study |
| **JM7492** | *lys3+:ptdh1*:NLS-linker-mTagBFP2:terminatortdh1:kanMX leu1Δ::Pcdc13::cdc13-L-cdc2as::cdc13 3'UTR::ura4+ cdc2Δ::kanMX6 cdc13Δ::natMX6 h+* | This Study |
| **JM7490** | *rga2Δ::kanMX6 lys3+:ptdh1*:NLS-linker-mTagBFP2:terminatortdh1:kanMX leu1Δ::Pcdc13::cdc13-L-cdc2as::cdc13 3'UTR::ura4+ cdc2Δ::kanMX6 cdc13Δ::natMX6* | This Study |
| **JM7502** | *rga4Δ::hphR lys3+:ptdh1*:NLS-linker-mTagBFP2:terminatortdh1:kanMX leu1Δ::Pcdc13::cdc13-L-cdc2as::cdc13 3'UTR::ura4+ cdc2Δ::kanMX6 cdc13Δ::natMX6* | This Study |
| **JM7479** | *lys3+:ptdh1*:NLS-linker-mTagBFP2:terminatortdh1:kanMX leu1-32:[pJK148-Pwee1-wee1-Twee1] h+* | This Study |
| **JM7558** | *lys3+:ptdh1*:NLS-linker-mTagBFP2:terminatortdh1:kanMX leu1-32:[pJK148-Pwee1-wee1-Twee1] rga4Δ::NAT* | This Study |
| **JM7532** | *rga2Δ::hphR lys3+:ptdh1*:NLS-linker-mTagBFP2:terminatortdh1:kanMX leu1-32:[pJK148-Pwee1-wee1-Twee1] h+* | This Study |
| **JM7503** | *ppa2Δ::kanMX6 lys3+:ptdh1*:NLS-linker-mTagBFP2:terminatortdh1:kanMX h+* | This Study |
| **JM7512** | *rga4Δ::hphR ppa2Δ::kanMX6 lys3+:ptdh1*:NLS-linker-mTagBFP2:terminatortdh1:kanMX* | This Study |
| **JM7533** | *rga2Δ::hphR ppa2Δ::kanMX6 lys3+:ptdh1*:NLS-linker-mTagBFP2:terminatortdh1:kanMX h+* | This Study |
| **JM7690** | *Diploid lys3+:ptdh1*:NLS-linker-mTagBFP2:terminatortdh1:kanMX / pil1-mCherry::natR* | This Study |
| **JM7674** | *Diploid rga2Δ::NAT lys3+:ptdh1*:NLS-linker-mTagBFP2:terminatortdh1:kanMX / rga2Δ::hphR lys3+:ptdh1*:NLS-linker-mTagBFP2:terminatortdh1:kanMX* | This Study |
| **JM7630** | *cdr2Δ::NAT zfs1Δ::kanMX6 ppa2Δ::kanMX6 lys3+:ptdh1*:NLS-linker-mTagBFP2:terminatortdh1:kanMX* | This Study |
| **JM7645** | *cdr2Δ::NAT rga4Δ::hphR zfs1Δ::kanMX6 ppa2Δ::kanMX6 lys3+:ptdh1*:NLS-linker-mTagBFP2:terminatortdh1:kanMX* | This Study |
| | *cdr2Δ::NAT rga2Δ::hphR zfs1Δ::kanMX6 ppa2Δ::kanMX6 lys3+:ptdh1*:NLS-linker-mTagBFP2:terminatortdh1:kanMX* | This Study |
| **JM7627** | *ppa2Δ::kanMX6 zfs1Δ::kanMX6* | This Study |
| **JM7381** | *cdr2Δ::NAT lys3+:ptdh1*:NLS-linker-mTagBFP2:terminatortdh1:kanMX* | Opalko *et al.,* 2022 |
| **JM6434** | *lys3+:ptdh1*:NLS-linker-mTagBFP2:terminatortdh1:kanMX* | Vještica et al., 2020 |

| | | |
|---|---|---|
| **JM6963** | *cdr2Δ::NAT rga4Δ::kanMX6 lys3+:ptdh1*:NLS-linker-mTagBFP2:terminatortdh1:kanMX cdc25-yomNeonGreen::hphR leu1:[pJK148-Pact-yomRuby3-term]* | This Study |
| **JM6831** | *rga2Δ::kanMX6 cdr2Δ::NAT lys3+:ptdh1*:NLS-linker-mTagBFP2:terminatortdh1:kanMX cdc25-yomNeonGreen::hphR leu1:[pJK148-Pact-yomRuby3-term] h-* | This Study |
| **JM6614** | *cdc13-mCherry-kanMX* | This Study |
| **JM7701** | *cdc13+-internal-sfGFPcp* | Kamenz et al., 2020 |
| **JM6713** | *cdc13-mNeonGreen-hphR h-* | This Study |
| **JM5290** | *cdc25-yomNeonGreen::hphR h+* | This Study |
| **JM656** | *sec8-1 ura4-D18 leu1-32 h+* | Lab collection |
| **JM7586** | *fim1-mNG::hphR h-* | This Study |
| **JM7588** | *arp2-mNG::mNG::hphR h-* | This Study |
| **JM5198** | *spn1-mNG::hphR h+* | This Study |
| **JM5288** | *mid1-yomNeonGreen::hphR h+* | This Study |
| **JM7587** | *cdc12-mNG::mNG::hphR h-* | This Study |
| **JM6907** | *cdc2-mNeonGreen-hphR lys3+:ptdh1*:NLS-linker-mTagBFP2:terminatortdh1:kanMX h-* | This Study |
